# Prefrontal and Subcortical Value Representation during Explore-Exploit Decision-Making and Suicide Attempts

**DOI:** 10.1101/2025.10.01.679761

**Authors:** Angela M. Ianni, Andrew Papale, Bea Langer, Aliona Tsypes, Katalin Szanto, Eran Eldar, Michael N. Hallquist, Alexandre Y. Dombrovski

**Affiliations:** Department of Psychiatry, University of Pittsburgh, Pittsburgh, PA, USA; Department of Psychology, University of North Carolina, Chapel Hill, NC, USA; Department of Psychology, Department of Cognitive and Brain Sciences, Hebrew University of Jerusalem, Jerusalem, Israel

## Abstract

**Background:** This study aims to understand learning and decision-making in suicidal behavior by investigating temporal dynamics of reward value encoding in individuals with late-life depression and a history of suicide attempts.

**Methods:** In a retrospective case-control study, 134 older adults (33 with depression and history of suicide attempts, 29 with depression and suicidal ideation but no past attempts, 32 with depression and no suicidal ideation/past attempts, and 40 psychiatrically healthy controls) completed an explore-exploit decision-making task during functional MRI (fMRI). Multilevel models of deconvolved fMRI time series, time-locked to trial events, interrogated whether temporal patterns of reward value encoding was associated with suicidal behavior.

**Results:** Suicidal behavior in general was associated with blunted ventral PFC (vPFC) value signals, but profiles varied as a function of attempt lethality. Specifically, low-lethality suicide attempts and excessive behavioral shifts were associated with abolished phasic responses to value updates in the default network vPFC and its connected regions, including the striatum, amygdala, and hippocampus. Additionally, an unexpected pattern of sustained negative value responses was observed in vPFC and striatal control subregions, and hippocampus of high-lethality suicide attempters.

**Conclusions:** Diverging patterns of decision-related responses may reflect different paths toward suicidal behavior. Impaired value updating in individuals with low-lethality suicide attempts suggests a failure to integrate recent outcomes alongside prior experience, potentially relating to over-reactivity to stressors and a lower threshold for suicide attempts. In contrast, increased control network responses to difficult choices in high-lethality attempters may underlie cognitive constriction and consideration of a narrow set of potential solutions.

## Introduction

Suicidal behavior becomes more lethal and resolute in late life (1,2), potentially in response to the burdens of aging. Yet, despite these burdens, suicide attempts remain relatively rare among older adults. Cognitive deficits (3–8), particularly those affecting decision-making, are among the key risk factors (9–13). Clinical theorists describe individuals in a suicidal crisis as cognitively overwhelmed (14), which prevents them from considering a broad range of options, including constructive alternatives to suicide. This narrowing of options can be understood through the lens of reinforcement learning (RL), a framework that explains how people adjust their expectations of success or the perceived value of choices based on feedback, and how they navigate the trade-off between trying new strategies and relying on familiar ones.

Our previous studies linked disrupted expected value representations in the ventral prefrontal cortex (vPFC) with impulsivity and poorly planned suicide attempts (11,15). People prone to suicidal behavior also display difficulties exploring a large set of options under time pressure (16). Specifically, individuals with a history of high-lethality suicide attempts demonstrated reduced exploration and often failed to shift away from unrewarded choices (“lose-shift”), suggesting a broader difficulty in going beyond a limited set of ineffective solutions. In contrast, low-lethality attempters tended to shift their responses excessively, potentially reflecting a tendency to abandon a course of action prematurely. The present study considers the neural substrates of these distinct profiles.

To resolve the explore-exploit dilemma, one balances known rewards against the potential of uncertain alternatives. This process involves maintaining and updating reward value expectations. Exploration is encouraged when the expected value of the best known option (“V_max_”) is low, incentivizing the discovery of better alternatives. Humans and other primates rely on the striatum and amygdala to resolve this dilemma when faced with a few discrete options (17,18). However, in more complex environments, mammals depend on hippocampal cognitive maps and flexible reward and goal representations in the vPFC (19–25).

Electrophysiological studies reveal the critical role of offline neural activity (i.e., intertrial activity) in integrating spatial information from the hippocampus with vPFC value and goal representations during learning and memory consolidation (26,27). Examining the temporal dynamics of vPFC value encoding within and between trials during exploration/exploitation (19), we found that subregions of the vPFC within the default mode network (“vPFC DMN”) and limbic network (28) are particularly sensitive to V_max_ (29). Whereas prior imaging studies assumed that value responses peak in the decision period, DMN and limbic activity was greatest offline, with vPFC DMN-hippocampal synchronization mediating exploratory behavior.

This study relaxes the temporal assumptions of prior imaging studies, including our own research on value representations in attempted suicide (11,15), examining vPFC and subcortical value responses both online and offline, as individuals explore and exploit a continuous environment under time pressure. We hypothesized that suicide attempts would be linked to disruptions in offline value encoding within the vPFC DMN and connected striatal and limbic subregions. We further predicted that these disruptions would scale with lose-shift behavior, reflecting a failure to integrate reinforcement over longer periods. Additionally, this study investigated how value encoding varies with the lethality of previous suicide attempts. Predicted trial-wise V_max_ estimates were derived from our validated computational model of adaptive explore-exploit transition (30) and entered into multilevel analyses of deconvolved blood-oxygen-level dependent (BOLD) responses (19) to investigate the within-trial timecourses of value encoding in the vPFC, striatum, hippocampus, and amygdala.

## Materials and Methods

### Sample characteristics

We recruited 134 older adults aged 49-80 from the greater Pittsburgh community, comprising four groups: 33 individuals with depression who had made at least one suicide attempt (suicide attempters, SA), 29 with depression and suicidal ideation but no history of suicide attempts (suicide ideators, SI), 32 with depression but no suicidal ideation or past suicide attempts (depressed, D-NS), and 40 psychiatrically healthy controls (nonpsychiatric controls, C). Suicidal ideation was assessed using the Scale for Suicidal Ideation [SSI (31)]. This is a subsample of the 143 individuals described in Tsypes et al. (16) who passed fMRI quality control. Group characteristics are summarized in Table 1, eTable 1, and eMethods. Secondary analyses divided the SAs into high and low lethality groups (HL and LL, respectively; HL n=12, 4 males; LL n=21, 7 males), using a cut-off of 4 on the Beck Lethality Scale [BLS (32)], corresponding to suicide attempts that were medically serious enough to result in hospitalization and would have been fatal without medical attention (33). The Institutional Review Board at the University of Pittsburgh approved the study procedures. Subjects provided written informed consent after receiving a complete description of the study.

**Table 1:** Demographic, clinical, and cognitive characteristics. Final study groups, after 12 participants were excluded due to failing fMRI quality control (see eTable 1 for full sample characteristics). Numeric values indicate mean (standard deviation). Test statistics and p-values for group differences across all groups are included, with post-hoc t-test results specifying specific group comparisons that are significant at a threshold of p<0.05. Measurement tools: Premorbid IQ from Wechsler Test of Adult Reading (WTAR (73)), dementia rating scale (DRS (74)), executive interview score of executive functioning (EXIT 25 (75)), physical illness burden from the cumulative illness rating scale-geriatric (CIRS-G (76)), depression severity from the Hamilton rating scale for depression (77), hopelessness from the Beck hopelessness scale score (78)), suicidal ideation from the Beck scale for suicidal ideation (79), antidepressant exposure from the Antidepressant Treatment History Form that rates exposure over the past 6 months from 0-5 based on dose and time of the medication (80), and lifetime substance use and anxiety diagnosis from clinical interview and chart review.

| Characteristic | Study group |  |  |  | Group difference test statistic (df) | P value | Post-hoc t-test results |
| --- | --- | --- | --- | --- | --- | --- | --- |
|  | Nonpsychiatric Controls (C)<br>(n = 40) | Nonsuicidal Depressed (D-NS)<br>(n = 32) | Depressed with Suicidal Ideation (SI)<br>(n = 29) | Depressed Suicide Attempters (SA)<br>(n = 33) |  |  |  |
| Male sex, No (%) | 18 (45.0) | 17 (53.1) | 13 (44.8) | 11 (32.2) | $\chi^2(3) = 2.6$ | 0.452 | |
| Age, y (sd) | 63.2 (8.1) | 61.8 (6.8) | 61.8 (4.9) | 61.5 (6.6) | F(3) = 0.5 | 0.695 |  |
| Race (White/Black/Asian/Multi-Race) | 37/3/0/0 | 25/6/1/0 | 27/1/1/0 | 27/4/0/2 | $\chi^2(9) = 12.0$ | 0.215 | |
| Educational level, y (sd) | 16.3 (2.6) | 15.6 (2.4) | 15.4 (2.0) | 14.3 (2.6) | F(3) = 4.3 | 0.007 | SA < C & D-NS |
| Premorbid IQ estimate (sd) | 111.4 (9.8) | 107.2 (14.6) | 114.2 (10.2) | 103.5 (11.4) | F(3) = 5.1 | 0.002 | SA < C & SI<br>D-NS < SI |
| Dementia rating scale score | 138.1 (3.5) | 135.1 (5.5) | 137.2 (3.9) | 135.4 (3.9) | F(3) = 3.9 | 0.010 | SA < C<br>D-NS < C |
| Executive interview score (sd) | 5.2 (2.4) | 6.4 (3.5) | 5.8 (2.6) | 6.5 (3.3) | F(3) = 1.7 | 0.179 |  |
| Physical illness burden | 4.5 (3.1) | 8.8 (4.5) | 10.1 (4.8) | 8.6 (4.2) | F(3) = 12.5 | <0.001 | SA, SI, D-NS > C |
| Hamilton Rating Scale for Depression score | 2.1 (2.5) | 13.3 (4.5) | 16.1 (6.3) | 14.5 (7.7) | F(3) = 50.2 | <0.001 | SA, SI, D-NS > C<br>SI > D-NS |
| Beck Hopelessness Scale score | 1.2 (1.3) | 5.5 (4.7) | 12.0 (6.0) | 8.9 (6.7) | F(3) = 29.8 | <0.001 | SA > C & D-NS<br>SI > C & SA |
| Suicidal ideation (Beck SSI) | 0 (0) | 3.3 (4.8) | 21.9 (4.5) | 27.8 (6.3) | F(3) = 322.6 | <0.001 | SA > C, D-NS, SI<br>SI > C, D-NS<br>D-NS > C |
| Antidepressant exposure, previous 6 months | NA | 21 | 22 | 28 | F(2) = 2.2 | 0.117 |  |
| Lifetime substance use, No. | NA | 14 | 14 | 23 | $\chi^2(2) = 5.0$ | 0.082 | |
| Lifetime anxiety, No. | NA | 16 | 19 | 27 | $\chi^2(2) = 7.3$ | 0.026 | SA > D-NS & SI |

### Explore-exploit behavior

Participants completed an explore-exploit decision-making task in a one-dimensional continuous space (“clock task”, Figure 1A, eMethods), where they decided when to stop a green dot revolving around a circle over a 5-second interval to earn rewards. They received probabilistic feedback controlled by two contingencies, such that unbeknownst to participants, expected values of choices either increased or decreased along the interval (Figure 1C), with reward probability and magnitude varying independently (eFigure 1A). The intertrial interval (ITI) ranged from 0.1 to 12.1 seconds (mean 4.1, standard deviation 3.5). Unsigned contingency reversals occurred after 40 trials and there were 240 total trials divided into two equal runs.

**Figure 1.**
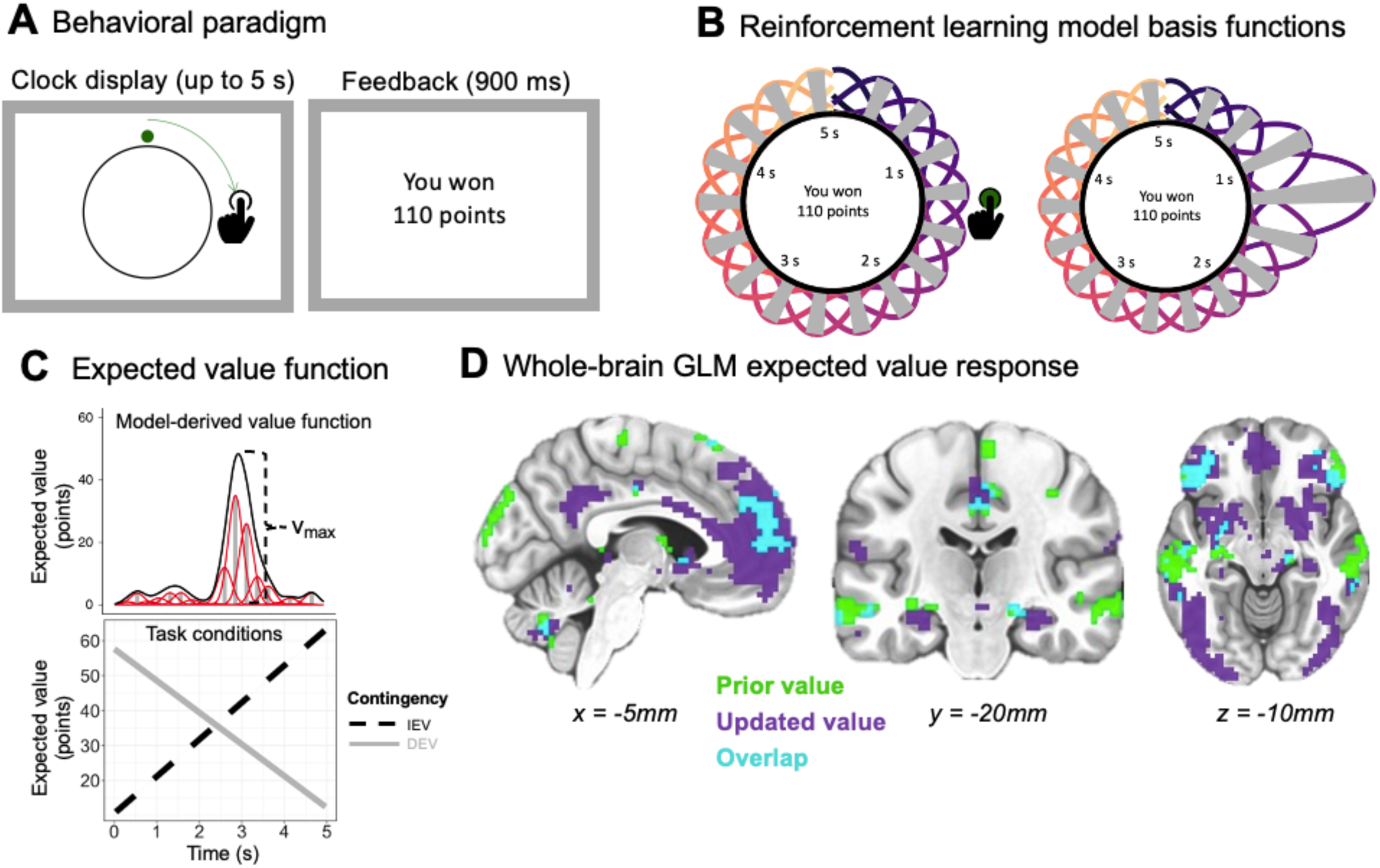
Paradigm and V_max_ signal. A) The clock paradigm consists of decision and feedback phases. During the decision phase, a dot revolves 360° around a central stimulus over the course of five seconds. Participants press a button to stop the revolution and receive a probabilistic outcome. B) Schematic of the SCEPTIC reinforcement learning model basis functions of expected value that tile the continuous space and are updated when feedback is received according to their proximity to the response (30). C) Linear representation of the expected value over the time interval after learning has occurred, with the maximum value (V_max_) highlighted (top). Rewards are drawn from one of two monotonically time-varying contingencies: increasing expected value (IEV) and decreasing expected value (DEV; bottom). D) Whole-brain GLM BOLD response to maximum value (V_max_), aligned to trial onset (prior value, green) and feedback (updated value, purple). Overlap between responses to prior and updated values is shown in cyan. Image is thresholded at p=0.005, one sided (z=2.6), unmasked, for visualization.

Participants were instructed that no points would be awarded if they failed to respond within the 5-second interval. Exploration of the interval manifested as shifts or swings in response time (“RT swings”), while exploitation was seen in choices that converge on the response time (RT) associated with V_max_. To enable assessment of individual differences in RT swing behavior, we extracted random slopes of RT swings following omissions for each individual using a linear mixed-effects model [lmer in R (34)] to predict RT in the current trial (RT_t_) from RT in the previous trial (RT_t-1_) (see eMethods for details). Conceptually, greater RT swings (i.e., larger shifts after unrewarded choices) would result in lower random slope values, as RT_t_ would differ more from RT_t-1_.

### Computational modeling of behavior

#### Core framework of the SCEPTIC RL Model

As illustrated in Figure 1B-C, eFigure 1B, and described in the eMethods, our model aimed to identify the highest-value region of the option space that a successful agent would exploit on each trial (V_max_), using each participant’s history of choices and reinforcement as the basis for estimation. To achieve this, we applied our validated StrategiC Exploration/Exploitation of Temporal Instrumental Contingencies (SCEPTIC) model (19,30,35), which approximates the expected value function along the 5-second interval using RL learning elements with selective maintenance and staggered receptive fields, implemented as Gaussian temporal radial basis functions (RBF; see model depiction in Figure 1B-C). Each basis function’s contribution to the integrated value representation is determined by its temporal receptive field Φ_!_

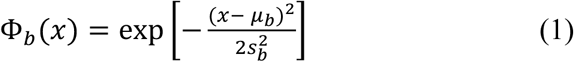

where *x* is an arbitrary point within the time interval *T*, and the mean and variance of the RBF are μ_*b*_ and *s*^2^_*b*_, respectively. Time-varying expected values are represented with a set of *B* weights, ***w*** = [*w*_1_, *w*_2_, …, *w_b_*], which scale the amplitudes of the corresponding basis functions to define the expected value function on trial *i*

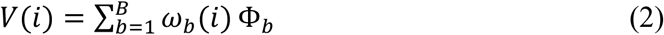

We placed the centers of 24 Gaussian RBFs evenly across the 5-second interval and set a fixed variance, *s^2^_b_*, so that each basis function overlapped 50% with its neighbor [for details and considerations of alternatives, see (36)]. The SCEPTIC model learns the expected values of different RTs by updating the weight of each basis function *b* according to the equation

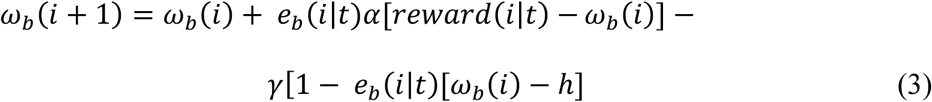

where *i* is the current trial on the task, *t* is the selected RT, and reward (*i* | *t*) is the reinforcement obtained on trial *i* given the choice *t*. Prediction error updates are weighted by the learning rate > and the temporal generalization function or eligibility *e*. Extending beyond traditional RL, the SCEPTIC model adds a selective maintenance parameter D between 0 and 1 that scales the degree of reversion toward a point *h*, taken to be 0 here. The eligibility *e_b_* of a basis function Φ_*b*_ to be updated by the prediction error is defined as the overlap with the temporal generalization function, *g*

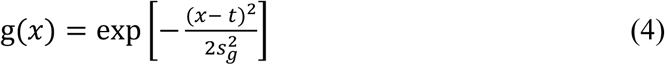

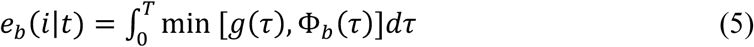

where τ represents an arbitrary time point along the interval *T*. For each RBF *b*, a scalar eligibility value *e_b_* (ranging from 0 to 1) quantifies the degree of overlap between the temporal generalization function and the RBF’s receptive field, where *e_b_* = 1 indicates complete overlap and *e_b_* = 0 indicates none. The basis functions are normalized to have a unit area under the curve, effectively treating them as probability density functions. Variance of the generalization function is fixed to match that of the basis functions (*s*^2^_*g*_ = *s*^2^_*b*_). Action selection in the SCEPTIC model is then governed by a softmax choice rule

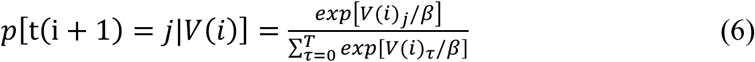

where *j* denotes a specific RT, and the temperature parameter β regulates value sensitivity, with higher β producing more stochastic and less value-driven choices. The SCEPTIC model outperformed two alternative approaches for this task—a traditional RL model without selective maintenance and a Kalman filter model incorporating parameters for value and uncertainty (see Supplemental Results)—as well as numerous other models in an independent sample (36).

#### SCEPTIC model parameter estimation

SCEPTIC model parameters were estimated from individual choices using an empirical Bayesian implementation of the variational Bayes approach (37). This method used a mixed-effects framework in which individual-level parameters were assumed to be drawn from a normally distributed population. Group-level summary statistics were inferred interatively by alternating between estimating population parameters and updating individual-level posteriors. Over successive iterations, individual priors were progressively shrunk towards the inferred population distribution, consistent with standard multilevel regression.

To minimize the possibility that individual differences in voxelwise estimates from model-based fMRI analyses reflected differences in parameter scaling, we refit the SCEPTIC model using the group-mean parameter values. This approach facilities comparison of regression coefficients across participants and reduces confounding in brain-behavior analyses due to variability in individual model fits. We extracted trial-wise V_max_ values for each individual to use as a parametric regressor in model-based fMRI analyses (30) (see eFig1B for example). Model parameters and fits were similar across groups (details in Supplemental results).

### fMRI analyses

3T fMRI data were collected during the clock task. Preprocessing details are in the eMethods. In addition to traditional whole-brain GLM analyses, timecourse analyses of offline, online, and immediate post-feedback responses were performed on deconvolved BOLD signal to functionally align the signal to the event of interest (19). After preprocessing, a leading hemodynamic deconvolution algorithm was applied voxelwise across the entire run to estimate neural population signals underlying the BOLD response (38). The deconvolved signal was extracted from -3 to +3 seconds around trial onset and from 0 to +3 seconds from feedback, then linearly interpolated onto a TR-based grid (0.6 seconds) centered around the event. We used multilevel regression models to compute trial-wise effects of V_max_ on the deconvolved signal from regions of interest in the vPFC and striatal DMN, limbic, and control networks (28), as well as the hippocampus and amygdala (see eMethods for additional details). We controlled for nuisance variables, including ITI, response time, outcome, unsigned prediction error, and age. False-discovery rate correction was applied in all analyses (39).

## Results

### Maximum value is encoded by medial PFC, striatum, and hippocampus

In our experiment, the same options (response times across the interval) are available on each trial, but their values are uncertain and must be learned from reinforcement. Accordingly, in whole-brain analyses we found V_max_ encoding in the medial PFC, striatum, and hippocampus, which was the strongest post-feedback (when value is updated), compared to trial onset (when no new information is revealed; Figure 1D, eTable 2). Group differences were evident only in trial onset-aligned analyses, with SA displaying blunted value signals in the striatum compared to SI (eFigures 2-3, eTable 3, and Supplemental results). However, these traditional GLM analyses treat offline activity between trials as an implicit baseline, precluding us from examining offline value responses and also complicating the interpretation of GLM results: it is impossible to know whether group differences reflect alterations in value modulation after events or offline, between trials. To address this, we used multilevel event-related deconvolved signal analysis (MEDuSA) to examine the temporal dynamics of value signals (19), as detailed in the next section.

### Timecourse of value responses in vPFC and subcortical networks. Effects of suicide attempt and lethality of attempt

Standard fMRI analysis approaches typically assume a fixed relationship between trial events (e.g., trial onset, choice, feedback) and the onset, height, and duration of the hemodynamic response function. MEDuSA avoides these assumptions by aligning deconvolved activity directly to trial events, as in electrophysiology, allowing observation of the full response shape. This approach is particularly useful for characterizing neural activity that unfolds offline, during the ITI. Note that MEDuSA does not overcome the temporal imprecision of BOLD response, leading us to interpret response peaks rather than apparent onsets and offsets.

In nonpsychiatric controls, V_max_ modulated offline activity in the vPFC default and limbic networks, as well as in the hippocampus, amygdala, and all striatal subregions [Figure 2A and 3A, prior value; see Papale et al., under revision, for replication in an independent sample (29)]. In striatal subregions receiving projections from limbic and control networks, high prior V_max_ predicted lower activity in the subsequent trial, consistent with reward prediction error dynamics (Figure 3A; see eFigure 4 for dynamics of BOLD response to reward and reward prediction error). In contrast, prior V_max_ responses persisted in the vPFC default network and hippocampus, matching axiomatic value signals. After feedback, updated value modulated activity in the vPFC default and limbic as well as in striatal default subregions (Figure 2A, 3A, “Updated value”). Critically, vPFC value modulation peaked during the offline period between trials.

**Figure 2.**
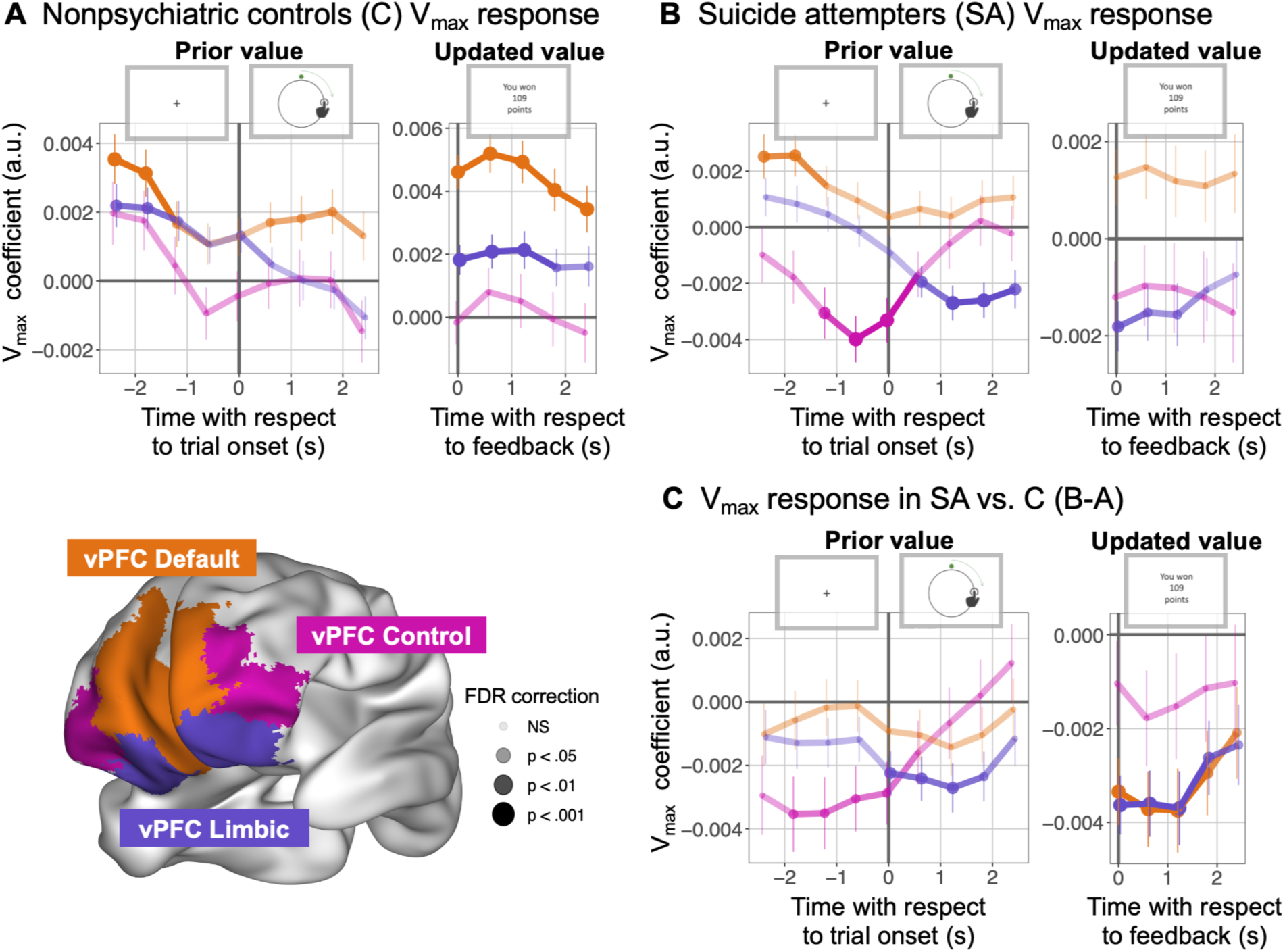
Dynamics of deconvolved BOLD V_max_ signal in the vPFC. A) vPFC responses to maximum value (V_max_) in nonpsychiatric controls (C) using multilevel analysis of deconvolved BOLD signal. Note the persistence of V_max_ encoding during the intertrial interval, which would be missed using traditional GLM approaches. B) vPFC V_max_ responses in suicide attempters (SA). C) Difference in vPFC V_max_ response between SA and controls (B-A). Repetition time (TR) is 0.6 seconds. Error bars denote standard errors from the multilevel model. Coefficients on the y-axis are in arbitrary units (a.u.). FDR correction was applied within the region (vPFC prior value=27 test statistics, vPFC updated value=15 test statistics), with significance indicated with circle size and opacity, defined in the figure legend.

**Figure 3.**
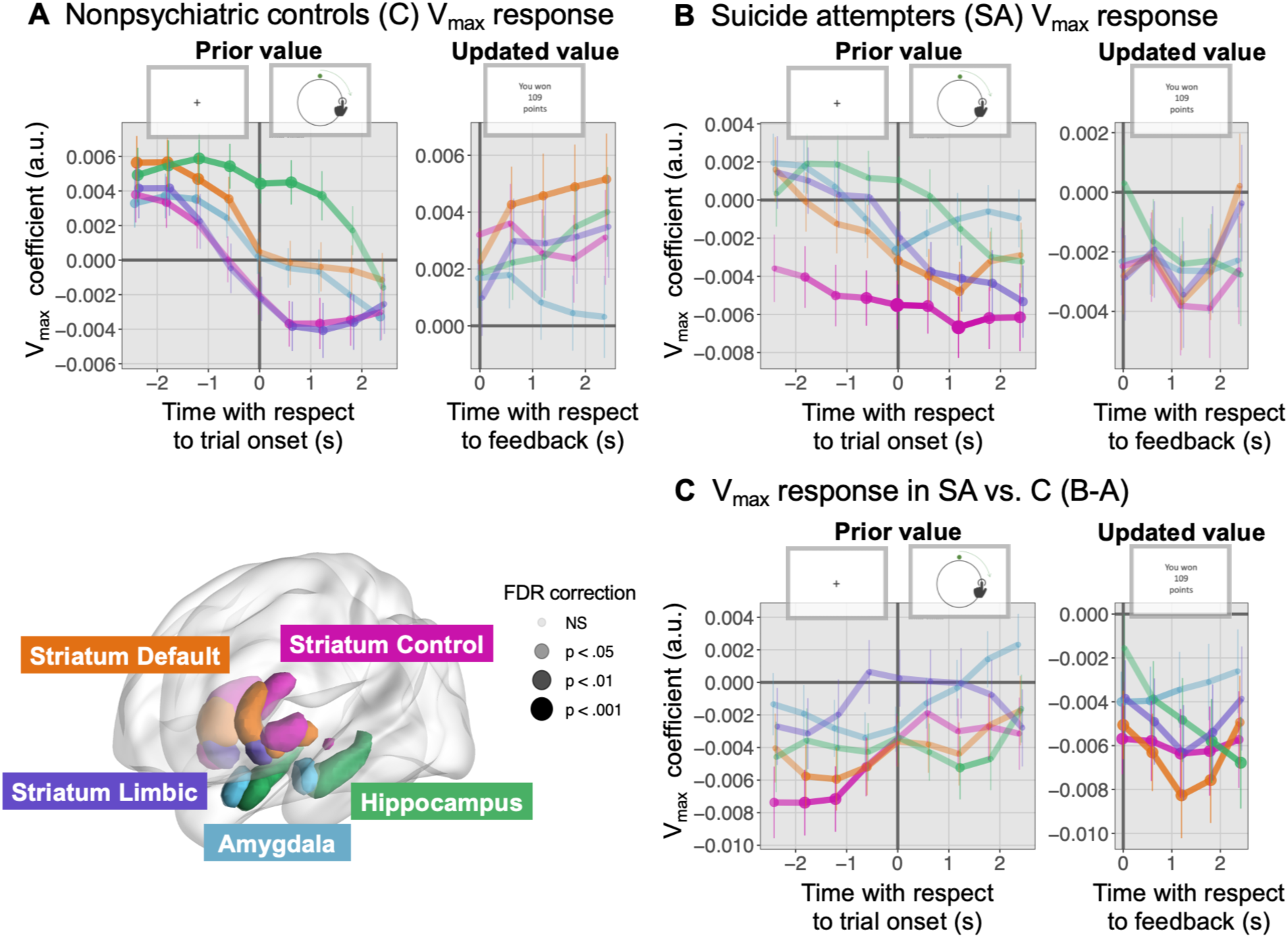
Dynamics of deconvolved BOLD V_max_ signal in the striatum, hippocampus, and amygdala. A) Subcortical responses to maximum value (V_max_) in nonpsychiatric controls (C), multilevel analysis of deconvolved BOLD signal. B) vPFC V_max_ responses in suicide attempters (SA). C) Difference between subcortical V_max_ response in SA compared to controls (B-A). Striatum subregions included in the ROIs are (1) default: head of the caudate nucleus, (2) limbic: ventral striatum, and (3) control: lateral putamen and tail of the caudate nucleus. Repetition time (TR) is 0.6 seconds. Error bars denote standard errors from the multilevel model. Coefficients on the y-axis are in arbitrary units (a.u.). FDR correction was applied within the region (subcortical prior value=45 test statistics, subcortical updated value=25 test statistics), with significance indicated with circle size and opacity, defined in the figure legend.

To identify robust group differences, we only interpret patterns differentiating SA from comparison groups at multiple contiguous time points, after FDR correction. SA displayed blunted V_max_ responses in default and limbic prefrontal and striatal subregions (Figure 2C, 3C). This blunting, most evident when V_max_ was updated post-feedback, was not observed in depressed comparison groups (group V_max_ responses in eFigure 5; comparisons in Figure 2-3 and eFigure 6-7). Furthermore, offline, suicide attempters showed a negative relationship between V_max_ and activity in control network vPFC and striatum. That is, their control network responses were stronger when V_max_ was low, i.e. when the optimal response location was unclear and decisions were more difficult. A similar pattern emerged among SA in vPFC limbic network online after trial onset (Figure 2B-C, left). As we probed the specificity of group differences, they remained unchanged in sensitivity analyses controlling for demographic and cognitive characteristics (eFigure 8), and there were no group differences in overall BOLD response to trial events (eFigure 9) or in total points obtained (Supplemental results). Altogether, SAs V_max_ responses were shifted negatively, abolishing responses in the reward-sensitive default network and producing negative modulation of the control network generally sensitive to task difficulty.

Curiously, whereas vPFC default and limbic network responses to updated value were most blunted in low-lethality SAs (LL; Figure 4A), sustained negative vPFC control network responses to V_max_ were only seen in the high-lethality subgroup (HL; Figure 4C). While LL displayed blunted striatal, amygdala, and hippocampal updated value responses immediately post-feedback, HL again displayed prominent, sustained offline negative responses of the control striatal subregions and hippocampus to V_max_ (Figure 4B, D; eFigure 10 for direct comparison of HL and LL SA). In summary, whereas LL SA displayed blunted phasic value updates after feedback, HL SA demonstrated sustained aberrant value responses across all epochs of the trial.

**Figure 4.**
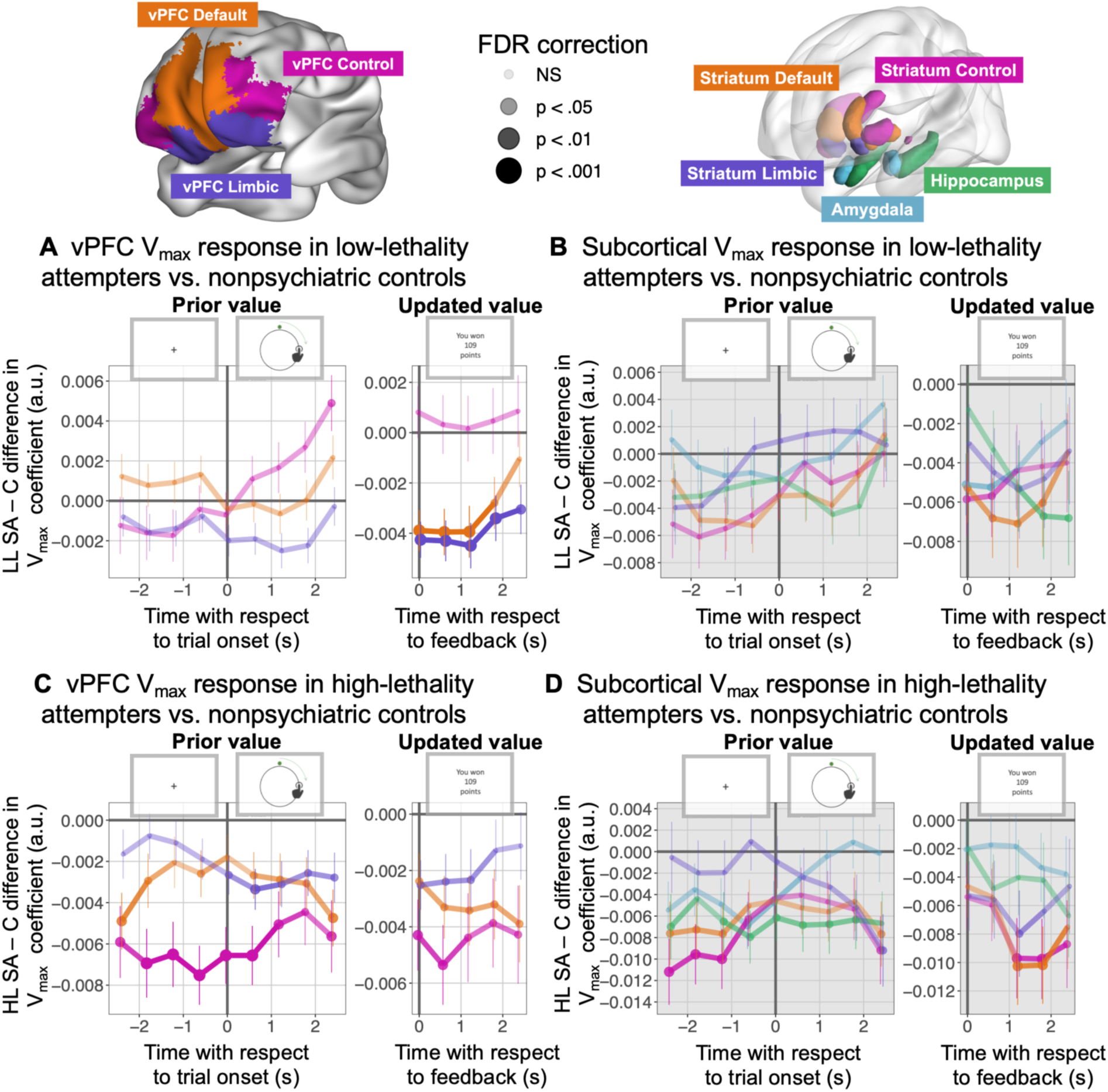
Value encoding deficits scale with suicide attempt lethality. Low-lethality suicide attempters (LL SA) have value encoding deficits specific to the post-feedback updating time period, while high-lethality attempters (HL SA) have broad value encoding deficits across all task phases. A) Difference in the vPFC V_max_ encoding between LL SA and nonpsychiatric controls (C). B) Difference in the subcortical V_max_ encoding between LL SA and controls. C) Difference in the vPFC V_max_ encoding between HL SA and controls. D) Difference in the subcortical V_max_ encoding between HL SA and controls. Striatum subregions included in the ROIs are (1) default: head of the caudate nucleus, (2) limbic: ventral striatum, and (3) control: lateral putamen and tail of the caudate nucleus. Repetition time (TR) is 0.6 seconds. Error bars denote standard errors from the multilevel model. Coefficients on the y-axis are in arbitrary units (a.u.). FDR correction applied within the region (vPFC prior value=27 points, vPFC updated value=15 points, subcortical prior value=45 points, subcortical updated value=25). Significance is indicated with circle size and opacity, defined in the figure legend.

### Lose-shifts are associated with blunted vPFC DMN value encoding

We reproduced our prior findings of limited behavioral exploration (decreased shifts from unrewarded choices) in HL vs. excessive choice shifts in LL [Figure 5B (16)]. The tendency to shift from unrewarded choices (random slopes of RT_t-1_ predicting RT_t_ for trials following omissions) was associated with diminished vPFC DMN V_max_ encoding at trial onset (GLM coefficients, Figure 5C). This correlation was significant in the entire sample and depressed individuals alone, and when using random slopes from all vs. only loss trials. Of note, while SA as a whole showed a trend toward diminished vPFC DMN V_max_ GLM coefficients at trial onset compared to C and SI (p_FDR_=0.058, eFigure 2A), there was no difference between the HL and LL SA groups (p_FDR_=0.983). There were no significant associations with value coefficients in any other vPFC or subcortical regions at trial onset or after feedback.

**Figure 5.**
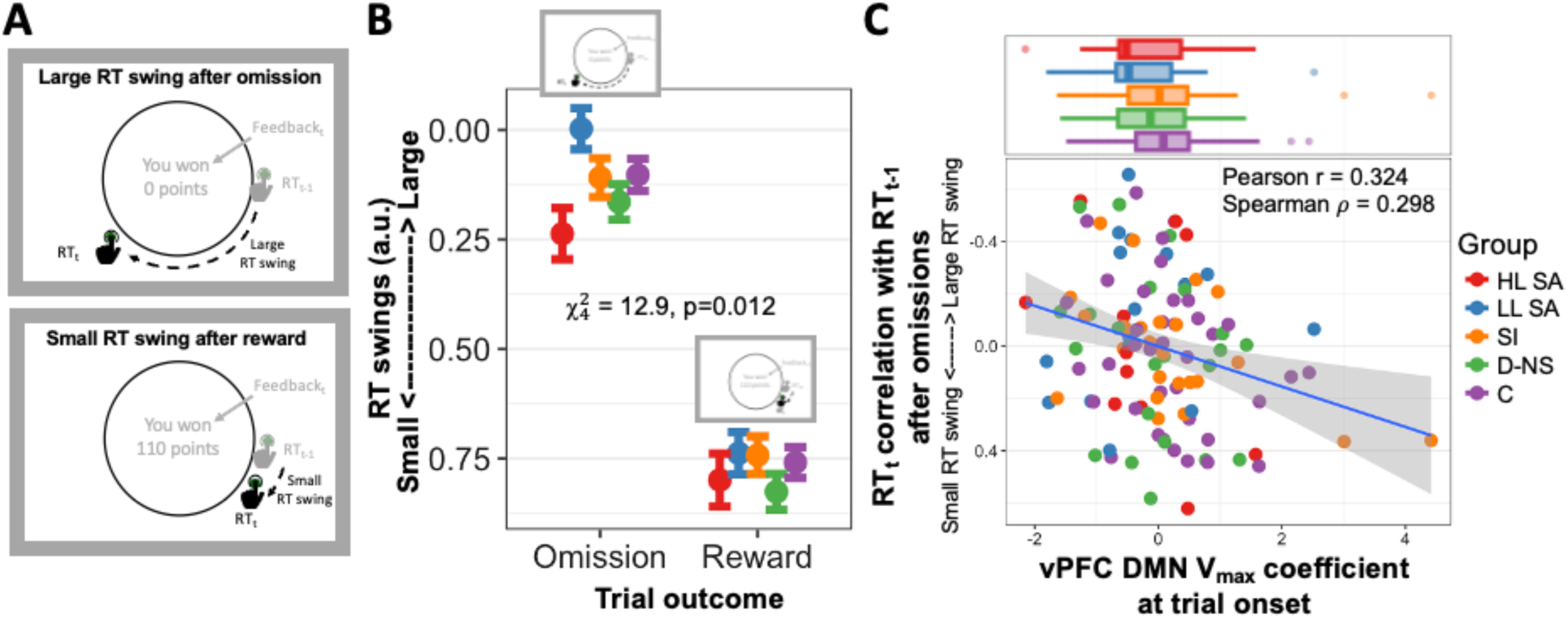
Exploratory response time (RT) swings are associated with vPFC DMN V_max_ encoding at trial onset. A) Schematic of RT swing behavior shows a large RT swing after an omission (top) and a small RT swing after a reward (bottom). B) Estimates from multilevel linear regression models predicting trial-level responses. We replicate group differences in RT swings in our fMRI sample (see Tsypes et al. 2024 for full sample results (16)), with HL SA displaying smaller RT swings and LL SA displaying increased RT swings after omissions. C) A greater tendency for RT swings after omissions is associated with diminished vPFC DMN value encoding at trial onset F_8,108_=2.51, p=0.015, vPFC DMN t=2.61, p=0.010; Pearson r = 0.324, p <0.001; Spearman rho=0.298, p=0.001). Coefficients on the x-axis were extracted from whole-brain GLM analyses aligned to trial onset with prior trial V_max_ as a parametric regressor. Box plots at the top of the figure display means, first and third quartiles (box), and 1.5*inter-quartile range (IQR, whiskers), with data outside of 1.5*IQR defined as outliers (circles). C = nonpsychiatric controls (purple), D-NS = nonsuicidal depressed (green), SI = depressed with suicidal ideation (orange), LL SA = low-lethality suicide attempters (blue), HL SA = high-lethality suicide attempters (red).

## Discussion

We replicated prior findings of reduced value-related signaling in the vPFC among individuals who had attempted suicide (11,15). Further, event-aligned analyses of BOLD signals revealed two distinct spatiotemporal patterns of aberrant neural responses. First, low-lethality suicide attempts were associated with abolished phasic default network and limbic responses to value updates in vPFC, connected striatal regions, amygdala, and hippocampus. Second, high-lethality attempts were associated with an unexpected pattern characterized by sustained negative value responses in vPFC and striatal control subregions and hippocampus. Below, we first consider our findings in the broader neurophysiological context and then discuss their behavioral and clinical relevance.

Individuals who had attempted suicide, particularly those with low-lethality attempts, showed blunted phasic value updates in the default/limbic vPFC and connected subcortical regions. This pattern was linked to excessive lose-shifting on the learning task (16), consistent with our main hypothesis. Lose-shift behavior is thought to depend on a rapidly decaying striatal memory trace (40,41) rather than on longer-term vPFC representations of reinforcement (42,43) and is normally suppressed by cognitive control (44). In this context, we interpret blunted value update signals as a failure to integrate recent reinforcement alongside the longer-term reinforcement history, resulting in choices driven solely by recent outcomes. Extrapolated to real life, these observations suggest a reduced awareness of the long-term consequences of one’s choices and heightened reactivity to recent experiences, which may lower the threshold for engaging in suicidal behavior. Further, as individuals attempt various solutions to their problems, they may prematurely abandon potentially beneficial strategies after a single setback.

We anticipated that behavioral heterogeneity among suicide attempters would map onto diverging neural alterations. However, whereas the pattern described above replicated our earlier findings (11,15), we also observed unexpected stronger responses to low-value options in the control vPFC and striatal subregions of suicide attempters (Figures 2B-C, 3B-C prior value and 4C-D), primarily those with high-lethality attempts. These responses occurred in granular cortex belonging to the frontoparietal control network: lateral orbital or ventrolateral prefrontal (vlPFC) area 11/47 and rostrolateral (rlPFC) areas 9 and 10. Both monkey and human studies implicate vlPFC in instrumental reversal learning and broader behavioral flexibility (45–47). Although the functions of rlPFC, which lacks clear monkey homologues, are less well understood, it may support relational category judgment (48,49), memory-based analogical reasoning (50), high-level sequential control (51,52) and, importantly, agency and strategic exploration (53,54). Of particular note is rlPFC’s sensitivity to the entire distribution of outcomes across the option set (54). Intriguingly, rlPFC lesions may even predict reduced suicidal ideation in traumatic brain injury patients (55). Notably, high-lethality suicide attempters, who tended to exhibit inflexible behavior and limited exploration on the learning task, showed stronger responses in the vlPFC and rlPFC during trials where the best option had a low value and the choices were more challenging. While this phenomenon requires replication and a fuller evaluation of its behavioral relevance, it may relate to narrow and ineffective problem-solving in a crisis and exclusive reliance on suicide over constructive alternatives.

Previous human fMRI studies, including our own, have generally assumed that vPFC primarily represents option values during online processing, either (i) when options are presented for a choice or (ii) when an outcome is delivered [reviewed in (23)]. However, human vPFC default network activity typically peaks during offline processing [(29,56–59), reviewed in (60)], and in monkeys and rats, vPFC value responses are at least as prominent offline as during choice or post-outcome (61–63). Consequently, we examined both offline and online value representations across vPFC networks. We found robust offline value encoding in default and limbic regions that weakened as participants started moving through the option space (Figure 2A, prior value) but increased after the value update (Figure 2A, updated value). Subcortically, ventral and dorsal striatal subregions—connected to limbic and control cortical networks, respectively—displayed prediction error responses that scaled inversely with prior value after feedback, as expected (64,65). Meanwhile, the hippocampus maintained sustained value signals, consistent with its role in integrating reinforcement into cognitive maps (19,66–71), with an intermediate pattern in striatal regions connected to the DMN. Altogether, these diverse spatiotemporal dynamics challenge the trial-aligned timecourse assumptions of conventional fMRI analyses and underscore the need for assumption-free approaches, such as our multi-level analysis of deconvolved BOLD, which treats within-trial time as completely general (72). This helps us understand why specific spatiotemporal alterations in value responses discussed above were fully apparent only in our assumption-free analyses (Figures 2-4) and not in conventional GLM analyses (eFigures 2-3).

A key strength of our study is the four-group case-control design differentiating attempted suicide from depression and suicidal ideation, as well as the inclusion of high-lethality suicide attempters and careful assessment and analysis of likely confounds. Furthermore, multi-level analyses of deconvolved BOLD acquired with a short TR of 0.6 seconds enabled us to relax the assumptions about activity timecourses that limit the utility and validity of conventional voxelwise deconvolution analyses. Finally, computational modeling provided insights into value encoding beyond those afforded by design factors and conventional model-free analyses.

The retrospective case-control design and the correlational nature of fMRI restrict causal inferences. The observed pattern of increased responses in vPFC control regions requires replication and further evaluation using brain stimulation studies.

In summary, low-lethality suicide attempters show blunted value updates in the default network, whereas high-lethality attempters exhibit heightened control network activity when faced with low-value options. These differences likely reflect two distinct behavioral pathways leading to suicide. Further investigation of these pathways is necessary for advancing personalized treatment and suicide prevention efforts.

## Author Contributions

Conceptualization: AI, AD. Investigation: AD, AP, BL, AT, MH. Resources: AD. Formal analysis: AI, AP. Writing: AI, AD, EE, MH, AT. Project administration: AD, KS.

## Acknowledgements

This work was funded by R01MH10095 (MRI) and R01MH085651 (recruitment, clinical, and cognitive characterization) from the National Institute of Mental Health. This research was supported in part by the University of Pittsburgh Center for Research Computing through the resources provided. Specifically, this work used the H2P cluster, which is supported by NSF award number OAC-2117681 and the HTC cluster, which is supported by NIH award number S10OD028483.

