## Supplementary Materials for "Prefrontal and Subcortical Value Representation during Explore-Exploit Decision-Making and Suicide Attempts"

### Supplemental material

*Ianni et al, 2025.*

#### Table of contents

##### Supplemental figures

eFigure 1: Task design and evolution of $V_{max}$ (p. 2)
eFigure 2. Group comparisons of BOLD $V_{max}$ extracted coefficients (p. 3)
eFigure 3: Exploratory whole-brain GLM $V_{max}$ group comparisons (p. 4)
eFigure 4. Response to reward and reward prediction error (p. 5)
eFigure 5. Individual group deconvolved neural responses to $V_{max}$ (p. 6)
eFigure 6. Dynamics of deconvolved BOLD $V_{max}$ signal in SA vs depressed control groups (p. 7)
eFigure 7. Dynamics of deconvolved BOLD $V_{max}$ signal in depressed control groups compared to non-depressed control group (p. 8)
eFigure 8. Sensitivity analysis – dynamics of value function timecourse and group comparisons, controlling for demographic characteristics and cognitive performance (p. 9)
eFigure 9. Group comparison of response to within-trial events (clock onset and feedback, intercept of activity independent of behavioral variables) (p. 10)
eFigure 10. Comparison of $V_{max}$ response in high-lethality and low-lethality suicide attempters (p. 11)

##### Supplemental tables

eTable 1: Demographic, clinical, and cognitive characteristics of complete sample (p. 12)
eTable 2. Whole-brain GLM voxelwise results for the parametric effect of $V_{max}$ (p. 13)
eTable 3. Whole-brain GLM group comparisons of $V_{max}$ response (p. 13)

##### eMethods

Participants (p. 14)
Mixed effects model of explore-exploit behavior (p. 14)
fMRI analyses (p. 15)
Acquisition and preprocessing (p. 15)
Voxelwise fMRI GLM analyses (p. 15)
Analyses of within-trial BOLD responses using voxelwise deconvolution (p. 16)

##### Supplemental results

Model validation across groups (p. 17)
Model-free behavioral results (p. 17)
Exploratory whole-brain analyses of group differences (p. 18)
Region of interest (ROI) fMRI GLM coefficient results (p. 18)
Timecourse fMRI analysis of reward and prediction error (p. 19)

##### References (p. 20)

### Supplemental figures

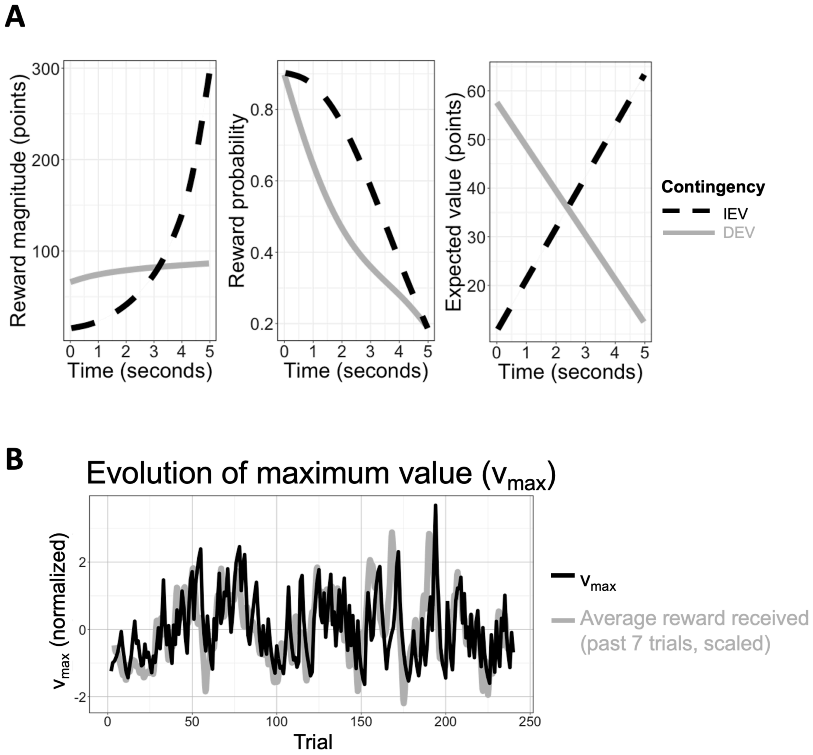


**eFigure 1: Task design and evolution of $V_{max}$.** A) Rewards are drawn from one of two monotonically time-varying contingencies: increasing expected value (IEV) and decreasing expected value (DEV). Reward probabilities and magnitudes vary independently. B) Model-derived estimated value maximum ($V_{max}$) over the experiment and the moving average reward received (over the past 7 trials, scaled) for an example participant.

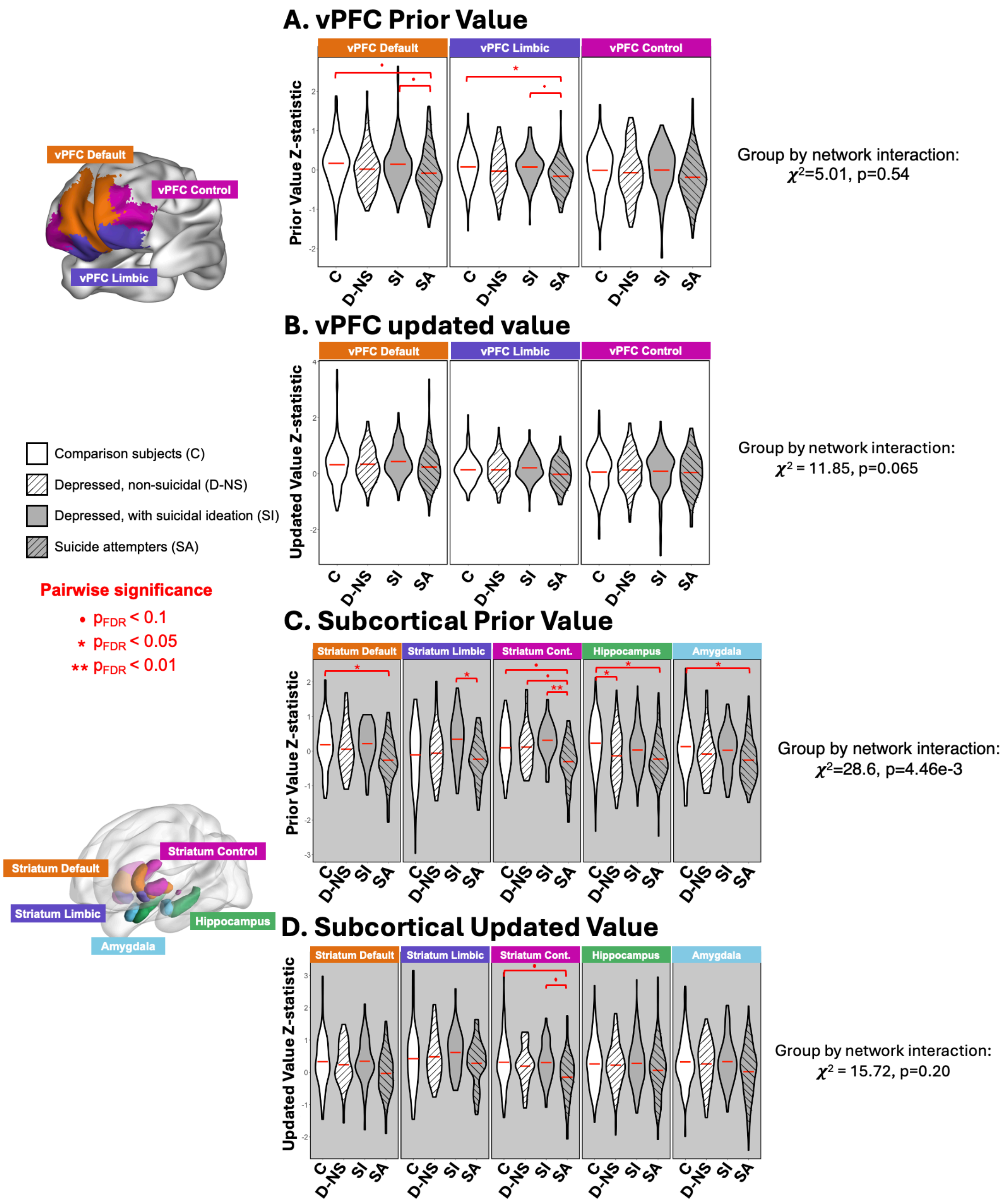


**eFigure 2. Group comparisons of BOLD $V_{max}$ extracted coefficients.** A) Group comparisons of extracted coefficients of prior value $V_{max}$ encoding in the default, limbic, and control networks regions of the ventral prefrontal cortex (vPFC). B) Group comparisons of extracted coefficients of updated value $V_{max}$ encoding in the vPFC. C) Group comparisons of extracted coefficients of prior value $V_{max}$ encoding in subcortical regions associated with the default, limbic, and control networks, the hippocampus, and the amygdala. D) Group comparisons of extracted coefficients of updated value $V_{max}$ encoding in subcortical regions. Striatum subregions included in the ROIs are (1) default: head of the caudate nucleus, (2) limbic: ventral striatum, and (3) control: lateral putamen and tail of the caudate nucleus.

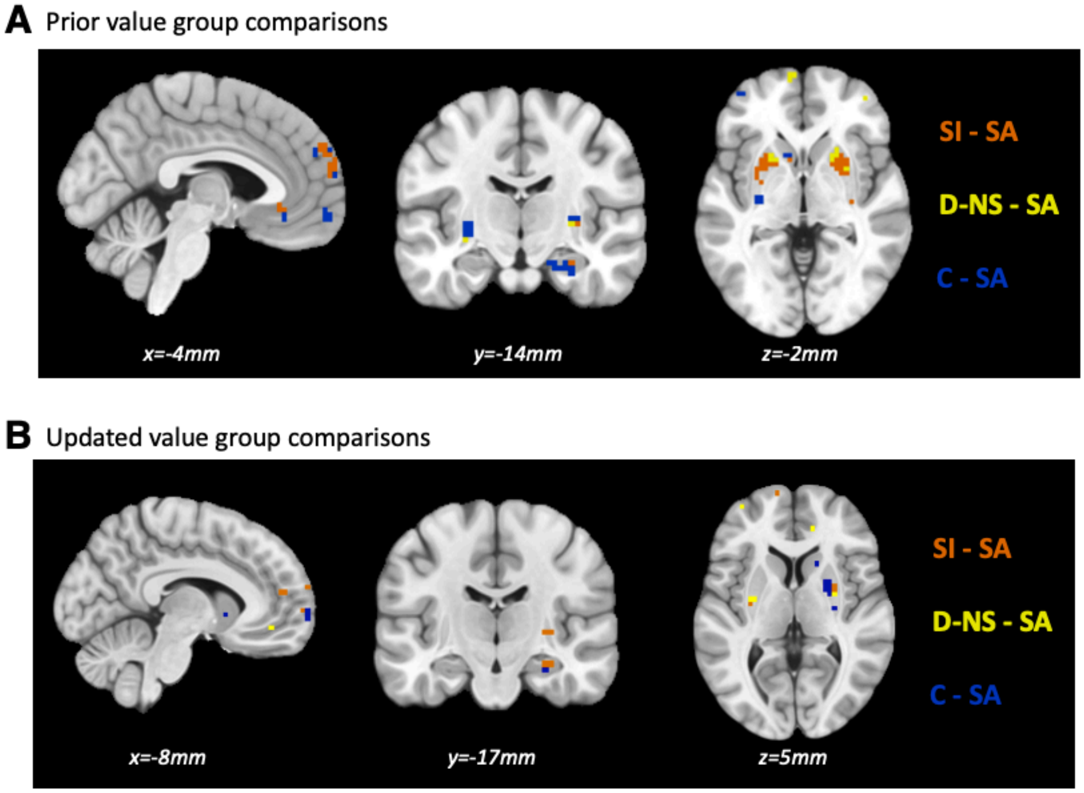


**eFigure 3: Exploratory whole-brain GLM $V_{max}$ group comparisons.** A) Prior value (aligned to trial onset) group comparisons of $V_{max}$. B) Updated value (aligned to feedback) group comparisons of $V_{max}$. Images are masked by ventral prefrontal cortex (vPFC), striatum, and hippocampus, and thresholded at $p<0.005$, one-tailed ($z=2.6$), uncorrected.

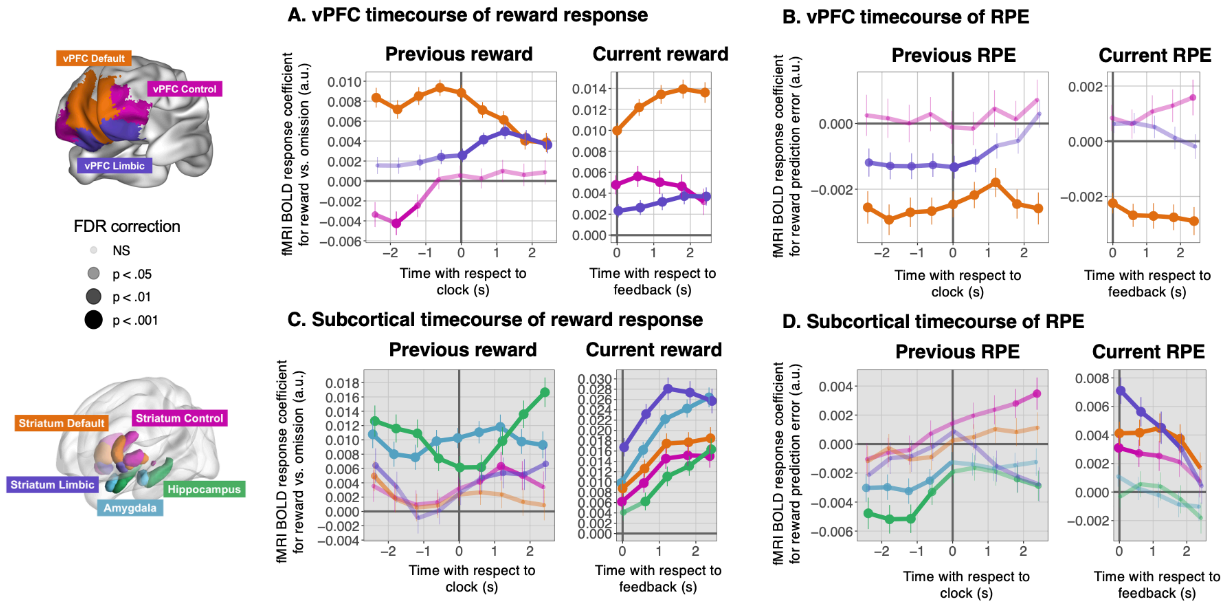


**eFigure 4. Response to reward (compared to omission) and reward prediction error (RPE, unsigned) in nonpsychiatric controls.** A) Timecourse of reward response to previous and current reward in the vPFC. B) Timecourse of previous and current unsigned RPE (absolute value) in the vPFC. C) Timecourse of reward response to previous and current reward in the subcortical regions. D) Timecourse of reward response to previous and current unsigned RPE in the subcortical regions. Striatum subregions included in the ROIs are (1) default: head of the caudate nucleus, (2) limbic: ventral striatum, and (3) control: lateral putamen and tail of the caudate nucleus. Repetition time (TR) is 0.6 seconds. Error bars denote standard errors from the multilevel model. Coefficients on the y-axis are in arbitrary units (a.u.). FDR correction was applied within the region (vPFC previous reward/RPE=27 test statistics, vPFC current reward/RPE=15 test statistics), with significance indicated with circle size and opacity, defined in the figure legend.

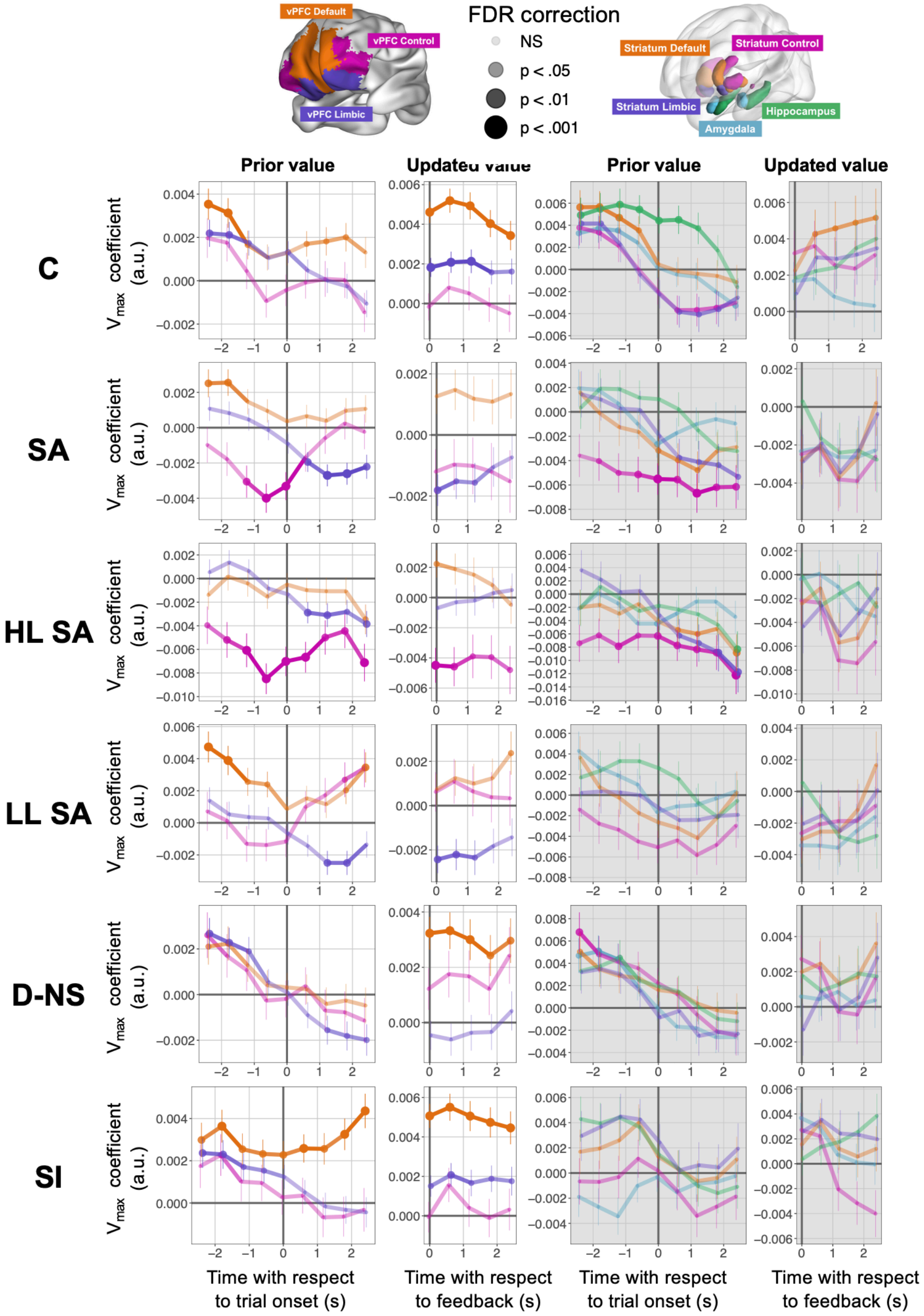


**eFigure 5. Individual group deconvolved neural responses to $V_{max}$.** Striatum subregions included in the ROIs are (1) default: head of the caudate nucleus, (2) limbic: ventral striatum, and (3) control: lateral putamen and tail of the caudate nucleus. Repetition time (TR) is 0.6 seconds. Error bars represent the SE of the estimate from the multilevel model. Coefficients on the y-axis are arbitrary units (a.u.). FDR correction applied within the region (vPFC prior value=27 points, vPFC updated value=15 points, subcortical prior value=45 points, subcortical updated value=25). Significance is indicated with circle size and opacity, defined in the figure legend. C = nonpsychiatric controls, SA = depressed suicide attempters, HL = high-lethality SA, LL = low-lethality SA, D-NS = nonsuicidal depressed, SI = depressed with suicidal ideation.

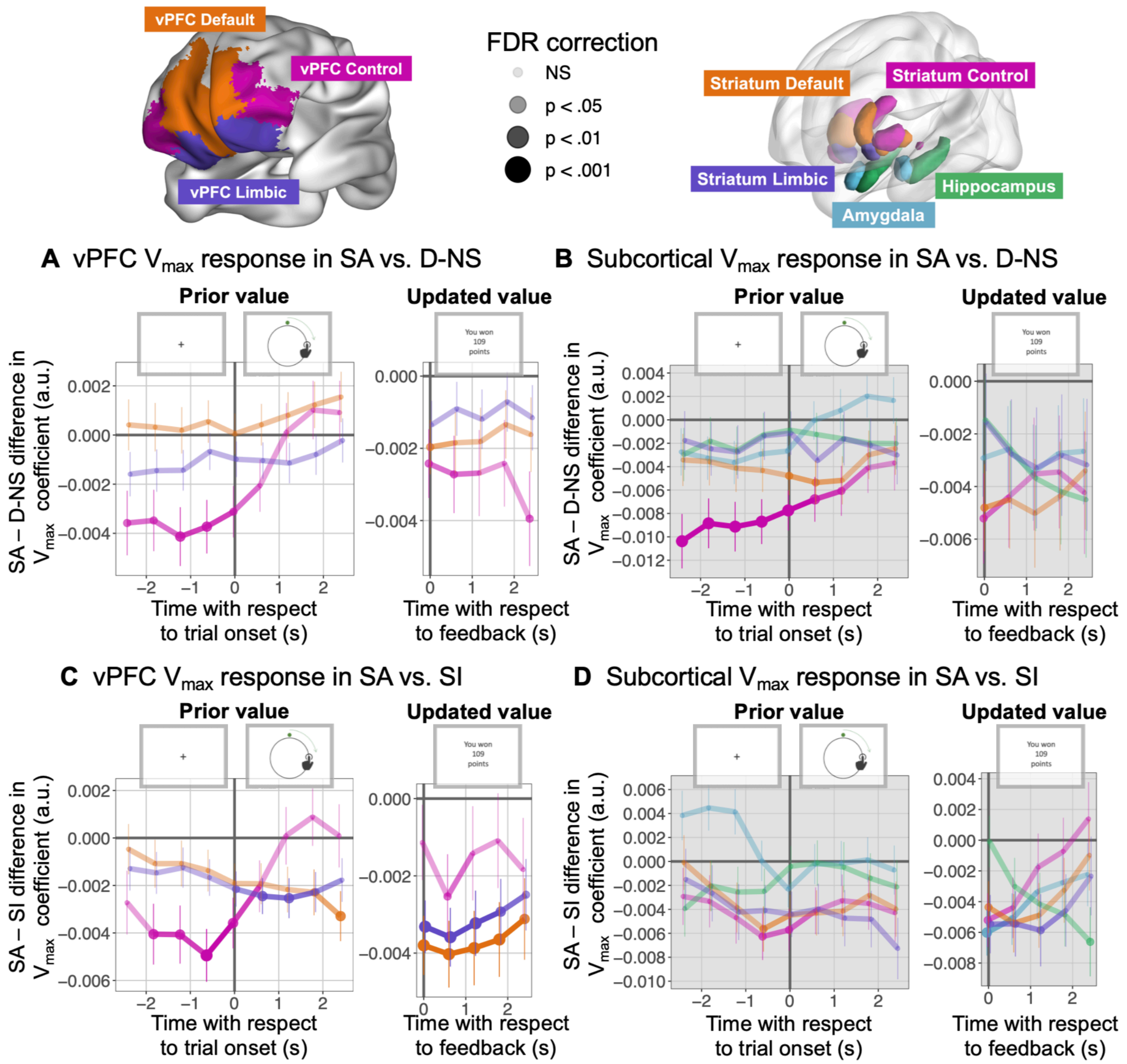


**eFigure 6. Dynamics of deconvolved BOLD $V_{max}$ signal in suicide attempters (SA) vs depressed control groups.** A) vPFC $V_{max}$ response in SA compared to D-NS. B) vPFC $V_{max}$ responses in SA compared to SI. C) Subcortical $V_{max}$ responses in SA compared D-NS. D) Subcortical $V_{max}$ responses in SA compared to SI. Striatum subregions included in the ROIs are (1) default: head of the caudate nucleus, (2) limbic: ventral striatum, and (3) control: lateral putamen and tail of the caudate nucleus. Repetition time (TR) is 0.6 seconds. Error bars represent the SE of the estimate from the multilevel model. Coefficients on the y-axis are arbitrary units (a.u.). FDR correction applied within the region (vPFC prior value=27 points, vPFC updated value=15 points, subcortical prior value=45 points, subcortical updated value=25), with significance indicated with circle size and opacity, defined in the figure legend. SA = depressed suicide attempters, D-NS = nonsuicidal depressed, SI = depressed with suicidal ideation.

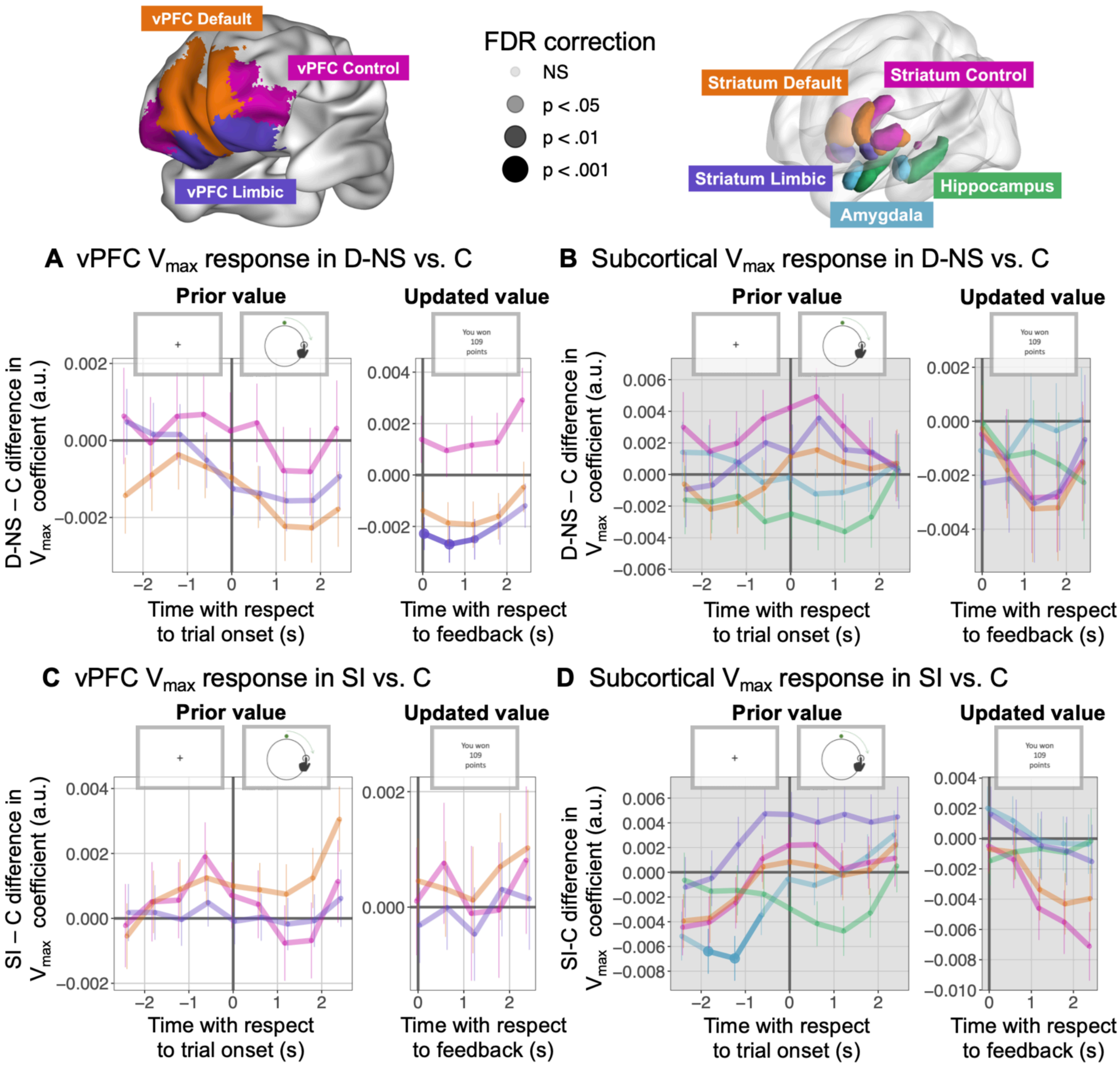


**eFigure 7. Dynamics of deconvolved BOLD $V_{max}$ signal in depressed control groups compared to non-depressed control group.** A) vPFC responses to maximum value ($V_{max}$) in non-suicidal depressed individuals compared to non-depressed comparisons subjects, multilevel analysis of deconvolved BOLD signal. B) vPFC $V_{max}$ responses in depressed individuals with suicidal ideation compared to non-depressed comparisons subjects. C) Subcortical $V_{max}$ responses in non-suicidal depressed individuals compared to non-depressed comparisons subjects. D) Subcortical $V_{max}$ responses in depressed individuals with suicidal ideation compared to non-depressed comparisons subjects. Striatum subregions included in the ROIs are (1) default: head of the caudate nucleus, (2) limbic: ventral striatum, and (3) control: lateral putamen and tail of the caudate nucleus. Repetition time (TR) is 0.6 seconds. Error bars represent the SE of the estimate from the multilevel model. Coefficients on the y-axis are arbitrary units (a.u.). FDR correction applied within the region (vPFC prior value=27 points, vPFC updated value=15 points, subcortical prior value=45 points, subcortical updated value=25), with significance indicated with circle size and opacity, defined in the figure legend. C = nonpsychiatric controls, D-NS = nonsuicidal depressed, SI = depressed with suicidal ideation.

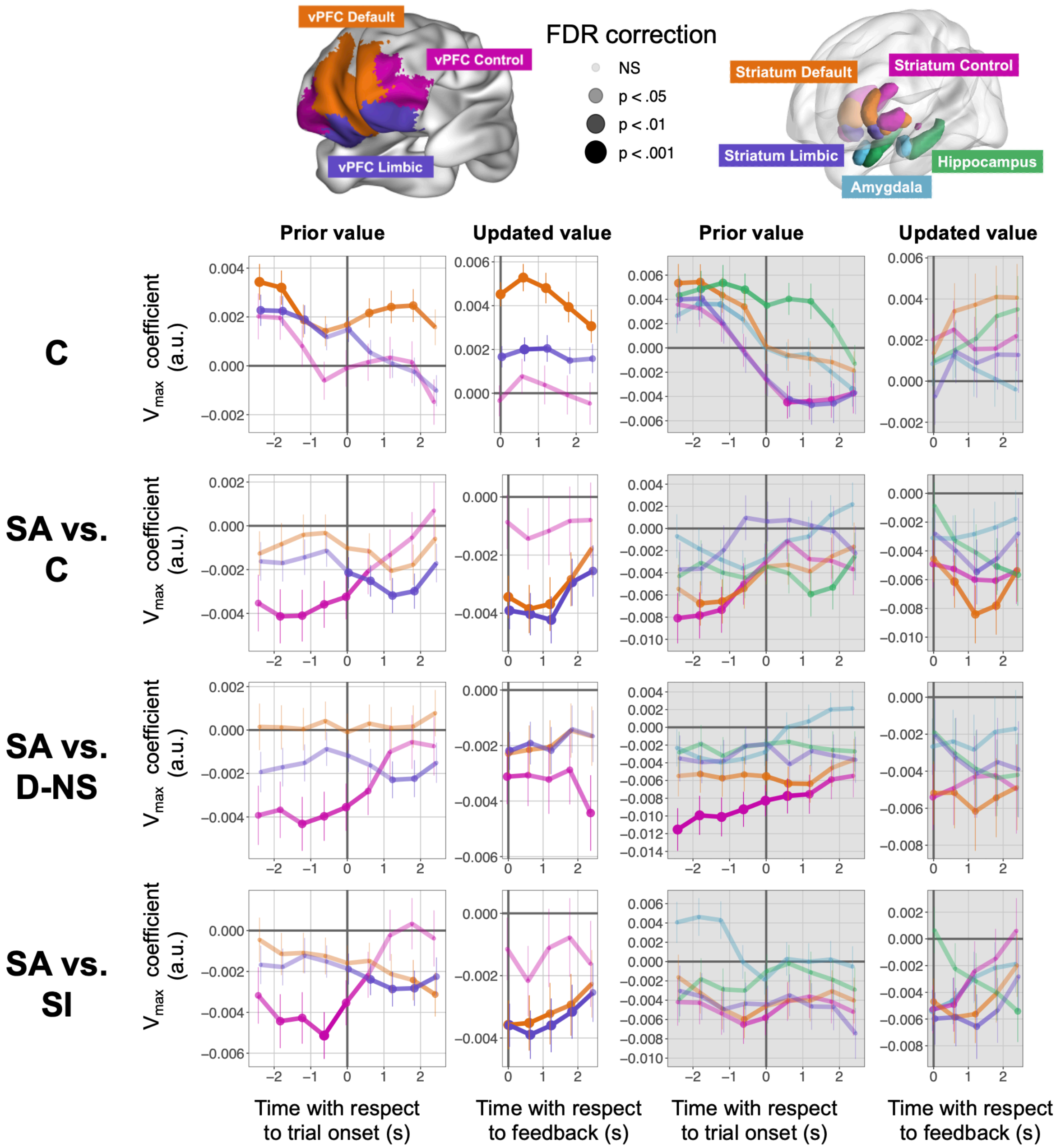


**eFigure 8. Sensitivity analysis – dynamics of value function timecourse and group comparisons, controlling for demographic characteristics and cognitive performance.** MEDuSAresults are qualitatively similar when adding additional demographic controls including IQ (WTAR), education years, dementia rating scale (DRS), burden of physical impairment (CIRSQ), depression (HAM-D), hopelessness (BHS), anxiety diagnosis, and antidepressant medication exposure. Striatum subregions included in the ROIs are (1) default: head of the caudate nucleus, (2) limbic: ventral striatum, and (3) control: lateral putamen and tail of the caudate nucleus. Repetition time (TR) is 0.6 seconds. Error bars represent the SE of the estimate from the multilevel model. Coefficients on the y-axis are arbitrary units (a.u.). FDR correction applied within the region (vPFC prior value=27 points, vPFC updated value=15 points, subcortical prior value=45 points, subcortical updated value=25), with significance indicated with circle size and opacity, defined in the figure legend. C = nonpsychiatric controls, SA = depressed suicide attempters, HL = high-lethality SA, LL = low-lethality SA, D-NS = nonsuicidal depressed, SI = depressed with suicidal ideation.

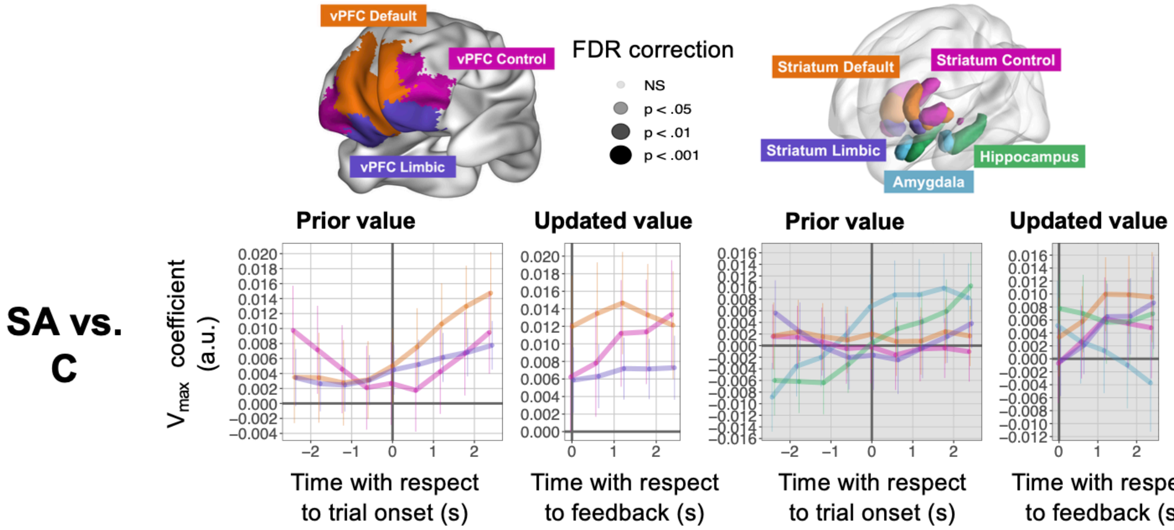


**eFigure 9. Group comparison of response to within-trial events (clock onset and feedback, intercept of activity independent of behavioral variables).** There were no significant differences between neural response to task events between suicide attempters and healthy comparison subjects. Striatum subregions included in the ROIs are (1) default: head of the caudate nucleus, (2) limbic: ventral striatum, and (3) control: lateral putamen and tail of the caudate nucleus. Repetition time (TR) is 0.6 seconds. Error bars represent the SE of the estimate from the multilevel model. Coefficients on the y-axis are arbitrary units (a.u.). FDR correction applied within the region (vPFC prior value=27 points, vPFC updated value=15 points, subcortical prior value=45 points, subcortical updated value=25), with significance indicated with circle size and opacity, defined in the figure legend. C = nonpsychiatric controls, SA = depressed suicide attempters.

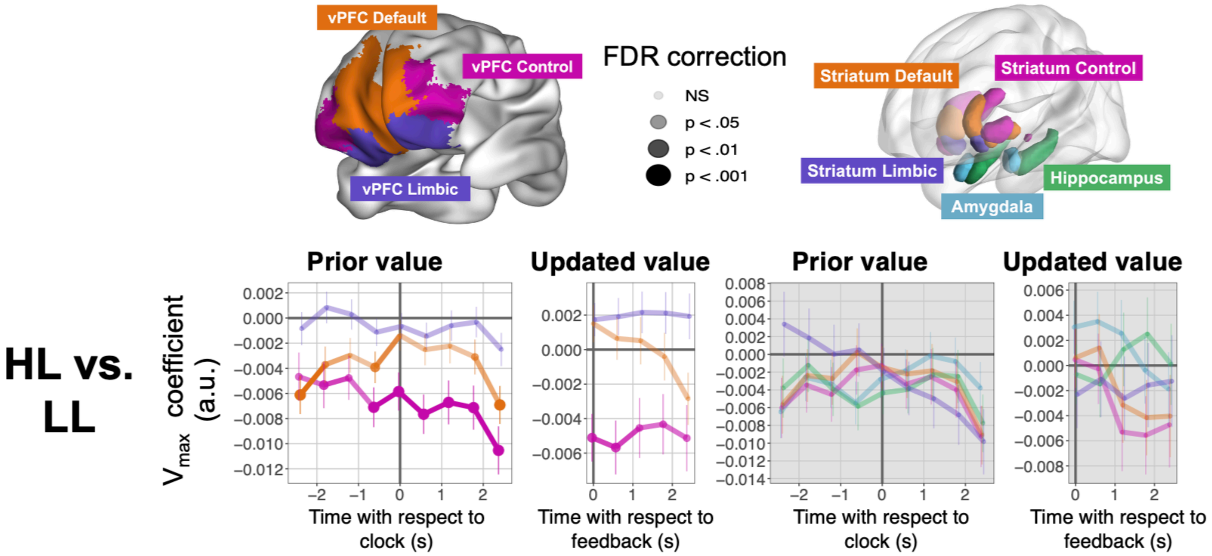


**eFigure 10. Comparison of $V_{max}$ response in high-lethality and low-lethality suicide attempters.** Striatum subregions included in the ROIs are (1) default: head of the caudate nucleus, (2) limbic: ventral striatum, and (3) control: lateral putamen and tail of the caudate nucleus. Repetition time (TR) is 0.6 seconds. Error bars represent the SE of the estimate from the multilevel model. Coefficients on the y-axis are arbitrary units (a.u.). FDR correction applied within the region (vPFC prior value=27 points, vPFC updated value=15 points, subcortical prior value=45 points, subcortical updated value=25), with significance indicated with circle size and opacity, defined in the figure legend. HL = high-lethality suicide attempters, LL = low-lethality suicide attempters.

### Supplemental tables

| Characteristic | Complete Study Group (with all participants including fMRI excludes) Nonpsychiatric Controls (C) (n = 43) | Nonsuicidal Depressed (D-NS) (n = 33) | Depressed with Suicidal Ideation (SI) (n = 33) | Depressed Suicide Attempters (SA) (n = 37) | Test statistic | P value | Post-hoc t-test results |
|---|---|---|---|---|---|---|---|
| Male sex, No (%) | 20 (46.5) | 18 (54.5) | 15 (45.5) | 12 (32.4) | $\chi^2 = 3.61$ | 0.307 | |
| Age, y (sd) | 63.3 (8.2) | 61.7 (6.7) | 61.6 (5.0) | 61.1 (6.5) | F = 0.82 | 0.484 | |
| Race (White/Black/Asian/Multi-Race) | 39/4/0/0 | 26/6/0/1 | 30/2/1/0 | 30/4/1/2 | $\chi^2 = 8.83$ | 0.453 | |
| Educational level, y (sd) | 16.5 (2.8) | 15.6 (2.4) | 15.3 (1.9) | 14.5 (2.7) | F = 4.40 | 0.005 | SA & SI < C |
| Premorbid IQ estimate (sd) | 111.7 (9.7) | 107.6 (14.5) | 112.8 (11.5) | 104.3 (11.2) | F = 3.91 | 0.010 | SA < C & SI |
| Dementia rating scale score | 138.1 (3.5) | 135.2 (5.5) | 137.1 (3.8) | 135.3 (4.1) | F = 4.15 | 0.007 | SA < C<br>D-NS < C |
| Executive interview score (sd) | 4.9 (2.5) | 6.3 (3.5) | 5.8 (2.7) | 6.4 (3.2) | F = 2.03 | 0.113 | |
| Physical illness burden | 4.6 (3.1) | 8.6 (4.5) | 9.7 (4.7) | 8.9 (4.5) | F = 11.56 | <0.001 | SA, SI, D-NS > C |
| Hamilton Rating Scale for Depression score | 2.0 (2.4) | 13.2 (4.4) | 15.8 (6.5) | 14.6 (7.7) | F = 53.36 | <0.001 | SA, SI, D-NS > C |
| Beck Hopelessness Scale score | 1.2 (1.3) | 5.9 (5.1) | 11.4 (6.0) | 8.5 (6.5) | F = 27.55 | <0.001 | SA > C, D-NS, SI<br>SI > C & D-NS<br>D-NS > C |
| Suicidal Ideation (Beck SSI) | 0.2 (1.5) | 3.2 (4.8) | 21.9 (4.2) | 27.7 (6.2) | F = 347.4 | <0.001 | SA > C, D-NS, SI<br>SI > C, D-NS<br>D-NS > C |
| Antidepressant exposure, previous 6 months | NA | 22 | 26 | 31 | F = 2.61 | 0.076 | |
| Lifetime substance use, No. | NA | 15 | 17 | 25 | $\chi^2 = 3.74$ | 0.154 | |
| Lifetime anxiety, No. | NA | 17 | 21 | 30 | $\chi^2 = 6.92$ | 0.031 | SA > D-NS & SI |

**eTable 1: Demographic, clinical, and cognitive characteristics of complete sample.** Complete study groups including the 12 excluded based on fMRI quality control. Numeric values indicate mean (standard deviation). Measurement tools: Premorbid IQ from Wechsler Test of Adult Reading (WTAR (1)), dementia rating scale (DRS (2)), executive interview score of executive functioning (EXIT 25 (3)), physical illness burden from the cumulative illness rating scale-geriatric (CIRS-G (4)), depression severity from the Hamilton rating scale for depression (5), hopelessness from the Beck hopelessness scale score (6)), suicidal ideation from the Beck scale for suicidal ideation (7), antidepressant exposure from the Antidepressant Treatment History Form that rates exposure over the past 6 months from 0-5 based on dose and time of the medication (8), and lifetime substance use and anxiety diagnosis from clinical interview and chart review.

| Analysis | Cluster size (# voxels) | Peak voxel MNI coordinates (mm) x | y | z | Peak-voxel z-stat | Location |
|---|---|---|---|---|---|---|
| Prior value | 191 | -62 | -45 | -1 | 6.615 | L middle temporal gyrus |
| | 166 | -59 | -61 | 27 | 6.836 | L angular gyrus |
| | 32 | -18 | -101 | 5 | 5.750 | L middle occipital gyrus |
| | 8 | 16 | -98 | 14 | 5.552 | R cuneus |
| | 7 | 28 | -79 | -38 | 5.373 | R cerebellum |
| Updated value | 6299 | 3 | 27 | 14 | 10.182 | bilateral anterior cingulate cortex, medial prefrontal cortex |
| | | -50 | 11 | 27 | 9.217 | L inferior frontal gyrus |
| | | 41 | 8 | 27 | 9.104 | R inferior frontal gyrus |
| | | -9 | 8 | -7 | 7.676 | L putamen/nucleus accumbens |
| | | -40 | 2 | 2 | 7.959 | L insula lobe |
| | | 16 | 8 | -10 | 7.723 | R putamen/nucleus accumbens |
| | | 41 | 2 | 2 | 7.220 | R insula lobe |
| | | 3 | -36 | 36 | 6.879 | Bilateral middle cingulate cortex |
| | | -3 | -51 | 30 | 6.996 | L posterior cingulate cortex |
| | | -25 | -20 | -17 | 6.603 | L hippocampus |
| | | 28 | -17 | -17 | 6.518 | R hippocampus |
| | 1517 | -47 | -51 | -20 | 8.99 | L inferior temporal gyrus |
| | | -50 | -54 | 24 | 7.437 | L angular gyrus |
| | | -59 | -42 | 30 | 7.339 | L supramarginal gyrus |
| | 366 | 50 | -79 | -1 | 7.624 | R inferior occipital gyrus |
| | | 44 | -58 | -20 | 7.110 | R fusiform gyrus |
| | 81 | 44 | -76 | -35 | 6.494 | R cerebellum |
| | 44 | -43 | -70 | -41 | 5.953 | L cerebellum |
| | 34 | -56 | -48 | 55 | 5.888 | L inferior parietal lobule |
| | 6 | -50 | -26 | -1 | 5.467 | L middle temporal gyrus |
| | 5 | -6 | -64 | -38 | 5.375 | L cerebellum |

**eTable 2. Whole-brain GLM voxelwise results for the parametric effect of $V_{max}$.** Clock-aligned (prior value) and feedback-aligned (updated value) clusters greater than 5 voxels, significant at a whole brain ptfce $p_{FWE}<0.05$ threshold. Results only include regions with positive value effects. Anatomical locations from Eickhoff-Zilles probabilistic cytoarchitectonic areas in MNI space (9).

| Analysis | Contrast | Peak voxel MNI coordinates (mm) x | y | z | Cluster size (# voxels) | Peak-voxel z-stat | Location |
|---|---|---|---|---|---|---|---|
| Prior value | Attempters < Ideators | -12 | 14 | -7 | 73 | 6.054 | L caudate nucleus |
| | | 22 | 8 | -1 | 18 | 5.364 | R putamen |
| | | 28 | 2 | -10 | 11 | 5.586 | |
| | | 28 | -14 | 5 | 10 | 5.431 | |
| | | 19 | -14 | 5 | 5 | 5.174 | |

**eTable 3. Whole-brain GLM group comparisons of $V_{max}$ response.** Results only include regions with positive value effects, which were found in trial onset-aligned (prior value) analyses. There were no significant group differences in feedback-aligned analyses. Statistics shown are for clusters meeting threshold of pTFCE pFWE<0.05, whole-brain bi-sided. Anatomical locations are from Eickhoff-Zilles probabilistic cytoarchitectonic areas in MNI space (9).

### eMethods

#### Participants

All study procedures received approval from the University of Pittsburgh Institutional Review Board (STUDY19030288). Before participating, all individuals gave written informed consent after being fully informed about the nature of the study. Participants were recruited from longitudinal studies drawing from inpatient and outpatient psychiatric services, as well as through advertisements in the greater Pittsburgh community. Eligible participants were between 50 and 80 years old at the time of enrollment. Exclusion criteria included any lifetime diagnosis of a psychotic or bipolar disorder, clinical signs of organic brain disease, medical conditions or treatments known to impact psychiatric functioning, and an IQ below 70 as measured by the Wechsler Test of Adult Reading (WTAR (1)). To better describe the sample, demographic details such as sex, race, and education level were collected via participant self-report using a structured list of options.

The group of depressed suicide attempters included individuals who had engaged in self-injurious behavior with intent to die within one month of the study assessments or who had made a past suicide attempt and were experiencing significant suicidal ideation at the time of enrollment. A psychiatrist confirmed each participant's history of suicide attempts using comprehensive sources including self-reports, medical records, and collateral information from the treatment team, family, and friends. Participants were excluded if there were major inconsistencies across these sources. The severity of suicide attempts was rated using the Beck Lethality Scale (BLS (10)), and for those with multiple suicide attempts, the attempt with the highest lethality score was considered. A BLS score of 4 or higher was used to classify high-lethality suicidal behavior. Suicidal ideation was measured using the Scale for Suicide Ideation (SSI (11)).

#### Mixed effects model of explore-exploit behavior

We used linear mixed effects modeling in R (lmer) used to test for group differences in exploratory behavior and to extract random slopes of RT swings following omissions. The full model is below:

$$(1)\ \mathrm{RT_t} \sim \mathrm{RT_{t-1}} * \mathrm{reward_{t-1}} * \mathrm{Group} + \mathrm{RT\_Vmax_{t-1}} * \frac{-1000}{\mathrm{trial}} * \mathrm{Group} + \mathrm{RT_{t-1}} * \frac{-1000}{\mathrm{trial}} * \mathrm{Group} + (1|\mathrm{id/run})$$

where response time (RT) is predicted by a linear combination of fixed effects including (1) the interaction of lagged response time from prior trial ($RT_{t-1}$), reward from prior trial ($reward_{t-1}$), and participant group, (2) the interaction of the response time with the maximum value from the previous trial ($RT\_Vmax_{t-1}$), the normalized negative inverse of trial in the run (-1000/trial), and group, and (3) the interaction of lagged response time, the normalized negative inverse of the trial in the run, and group. The model also includes random effects for individuals and run number.

#### fMRI analyses: Acquisition and preprocessing

fMRI imaging was conducted at the University of Pittsburgh's Magnetic Resonance Research Center using a Siemens Magnetom Prisma scanner. Functional images were acquired with a T2*-weighted EPI sequence optimized for fast TR (TR = 600ms, TE = 27ms, flip angle = 45°), using a multiband acceleration factor of 5 and a voxel size of 3.1 $mm^3$. High-resolution T1-weighted anatomical images were collected for coregistration (voxel size = 1.0 $mm^3$, TR = 2.3s, TE = 3.35ms, GRAPPA 2x acceleration). A gradient echo fieldmap (TE = 4.47ms and 6.93ms) was also acquired to correct for magnetic field inhomogeneity.

fMRI data were preprocessed using FMRIPREP version 20.1.1 (12), a Nipype-based tool (13). T1-weighted images were corrected for intensity non-uniformity using N4BiasFieldCorrection v2.1.0 (14) and skull-stripped with antsBrainExtraction.sh v2.1.0 using the OASIS template. Brain surfaces were reconstructed using FreeSurfer's recon-all v6.0.1 (15), and the brain mask was refined with a custom variation of the method to reconcile ANTs-derived and FreeSurfer-derived segmentations of the cortical gray-matter of Mindboggle (16). Spatial normalization to the ICBM 152 Nonlinear Asymmetrical template version 2009c (17) was performed using antsRegistration (ANTs v2.1.0, (18). Brain tissue was segmented into CSF, white matter, and gray matter using FSL FAST v5.0.9 (19).

Functional data were slice-time corrected using AFNI's 3dTshift v16.2.07 (20), and motion corrected with FSL's MCFLIRT (FSL v5.0.9 (21)). Fieldmap-based distortion correction was applied using FSL-s fugue (22), followed by coregistration to T1-weighted images using FreeSurfer's bbregister with boundary-based registration (six degrees of freedom (23)). All spatial transforms (motion correction, distortion correction, BOLD-to-T1-weighted, and T1-weighted to MNI) were combined and applied in a single resampling step using antsApplyTransforms (ANTs v2.1.0) with Lanczos interpolation.

Frame-wise displacement (24) was computed for each run using Nipype. To mitigate motion-related artifacts, we used ICA-AROMA (19–25)(25) to identify and regress out noise components via FSL's regfilt, using a non-aggressive denoising strategy. We then applied a temporal high-pass filter (cutoff = 0.008 Hz) to remove low-frequency drifts (26). The same filter was applied to all regressors in subsequent MEDuSA analyses. Finally, voxelwise time series were normalized to a mean of 100 to ensure comparability of regression coefficients across runs and participants.

#### Voxelwise fMRI GLM analyses

We conducted a whole-brain GLM analysis in FSL (27) to identify regions where fMRI BOLD responses were modulated by $V_{max}$. SCEPTIC-derived parametric modulators for $V_{max}$ were convolved with trial onset (reflecting prior value) or feedback (reflecting updated value), modelled as a conventional unit-height regressor. $V_{max}$ was mean centered within run to orthogonalize broad trial effects from specific $V_{max}$ responses. Group analyses controlled for age and were corrected for multiple comparisons using the Probabilistic Threshold-free Cluster Enhancement method (28) at $p < 0.05$. In addition, we extracted GLM coefficients for prior $V_{max}$ (aligned to trial onset) and updated $V_{max}$ (aligned to feedback) for each participant. Coefficients were extracted from atlas-defined functional networks (DMN, limbic, control networks of vPFC and striatum, hippocampus, and amygdala) (29) by averaging all voxels within each region of interest. We used linear regression (lm in R (30)) to test the effects of standardized value coefficients on RT swing behavior.

#### Analyses of within-trial BOLD responses using voxelwise deconvolution

We applied a leading hemodynamic deconvolution algorithm to estimate uneural activity from the BOLD signal (31). Voxelwise BOLD data were deconvolved for each subject, and the resulting time series were averaged within predefined cortical and subcortical parcels: specifically, the vPFC parcels corresponding to the default, limbic, and control networks, and subcortical regions including areas of the striatum connected to the default, limbic, and control networks as well as the hippocampus and amygdala. These averages were organized into a region x time matrix for each fMRI run.

To estimate trial-level neural responses, we extracted segments of the deconvolved signal aligned to task events, specifically, from -3 to +3 seconds around clock onset and from 0 to +3 seconds following feedback. We censored time points that overlapped with adjacent trials. To facilitate the application of discrete-time models, these event-aligned signals were resampled onto an evenly spaced temporal grid using linear interpolation, matching the repetition time (TR) of the scan (0.6 seconds). This interpolation resampled the data without altering its temporal resolution (i.e., no upsampling or downsampling was performed).

For each subject, this procedure yielded a trial x time point x region matrix: 240 trials x 11 or 6 time points (for clock- and feedback-aligned data, respectively) x 3 or 5 region (for vPFC and subcortical analyses, respectively). These matrices were concatenated across participants for group-level analysis. To model the relationship between neural activity and key decision variables, we performed multilevel regression analyses using the lme4 (32) package in R (v. 4.3.1). Each model included a random intercept for subject and was run separately for each time x region combination. Predictor variables included core decision-related measures (see equations 2 and 3), and sensitivity analyses additionally controlled for sex, premorbid IQ (wtar), years of education, dementia rating scale score, cumulative illness rating scale-geriatric score, Hamilton depression score, Beck hopelessness score, antidepressant medication exposure in the past 6 months, and categorical anxiety diagnosis. These regression coefficients provide estimates of when where specific decision-related signals, such as $V_{max}$, are reflected in event-aligned neural activity.

Clock-onset aligned:

$$(2)\ \text{deconvolved\_activity} \sim \text{Group} * \text{Vmax}_{t-1} + \text{ITI}_{t-1} + \text{RT}_{t-1} + \text{reward}_{t-1} + \text{abs(PEmax)}_{t-1} + \text{age} + (1|\text{id})$$

Feedback-aligned model:

$$(3)\ \text{deconvolved\_activity} \sim \text{Group} * \text{Vmax}_{t} + \text{ITI}_{t-1} + \text{ITI}_{t} + \text{RT}_{t-1} + \text{RT}_{t} + \text{reward}_{t-1} + \text{reward}_{t} + \text{abs(PEmax)}_{t} + \text{age} + (1|\text{id})$$

Importantly, due to the temporal smoothness of the BOLD signal, deconvolved time series remain highly autocorrelated, limting the precision of temporal inferences. Furthermore, this temporal (and potentially spatial) autocorrelation introduces dependencies across models, violating assumptions of independence. To correct for multiple comparisons under these conditions, we applied the Benjamini-Yekutieli correction across all terms of interest, controlling the false discovery rate (FDR) at 0.05 (33).

### Supplemental results

#### Model validation across groups

We modeled individuals' behavior using a previously validated reinforcement learning (RL) model, Strategic ExPloration/ExPloitation of Temporal Instrumental Contingencies (SCEPTIC) (34), which is a variation of a traditional RL model that includes a decay parameter such that unchosen values decay towards zero. We used the Variational Bayesian Analysis (VBA) toolbox for model-fitting (https://mbb-team.github.io/VBA-toolbox/). We first ran a mixed effects model that allows model parameters to vary between individuals. We compared the SCEPTIC model to two other models for a subset of 120 participants: (1) a traditional RL model without the decay parameter and (2) a Kalman filter model with parameters representing value and uncertainty. We found that, in accordance with prior studies in young healthy adults (34), the SCEPTIC model was the best fit for the data in this sample (all subjects: n =120, exceedance probability = 1, BOR = 6.79e-25, estimate model frequency = 0.741; C: n=27 exceedance probability = 0.99839, BOR = 1.1e-5, estimated model frequency = 0.761; D-NS: n=26, exceedance probability = 0.96754, BOR = 1.86e-4, estimated model frequency = 0.665; SI: n=29, exceedance probability = 0.99815, BOR = 1.73e-5, estimated model frequency = 0.747; SA: n=33, exceedance probability = 0.99827, BOR = 3.55e-6, estimated model frequency = 0.733). Model-fit, as measured by R-squared, did not differ between groups (F=2.026, p=0.11, df=3). The SCEPTIC model contains three free parameters, learning rate (alpha), temperature (beta), and the selective maintenance parameter (gamma) that decays unchosen values towards zero. Likewise, model parameter values were similar across groups (alpha: F=2.205, p=0.0902; beta: F=0.105, p=0.957; gamma: F=0.119, p=0.949, df=3). We then ran a fixed effects model using the group mean for each model parameter as the prior estimates and used the final parameter estimates across the group to generate parametric regressors to be used in the imaging analyses. In this study, we focus on the trial wise estimated value parametric regressor.

#### Model-free behavioral results

There were no group differences in total score obtained over the experiment (F=1.62, p=0.182, df=3). We used a multilevel model to investigate model-free behavior with response time as the independent variable and prior trial response time, outcome of prior trial, most valuable response time selected so far, trial, and group as dependent variables. We also included the interactions of group with prior trial response time by outcome of prior trial, most valuable response time by trial, and prior trial response time by trial as dependent variables. We allowed for random intercepts for individual and run levels. Overall, we found that response time was highly correlated with previous trial response time ($\chi^2$=970.46, p=4.74e-213, df=1) and outcome ($\chi^{2=}$61.36, p=4.76e-15, df=1). When deciding to explore, participants shift their response time away from the previous trial. Therefore, we expect that greater changes in response time compared to the previous trial are related to decisions to explore alternative options. Looking at all participants, we did find an interaction of previous trial outcome on the tendency to change response time ($\chi^2$=731.90, p=3.45e-161, df=1), but there was no difference when comparing between groups ($\chi^2$=93, p=0.82, df=3). However, as previously published, we did find an effect of suicide attempt lethality such that high lethality attempters shifted responses less after losses compared to other groups (35).

#### Exploratory whole-brain analyses of group differences

Given the negative inflection in neural-response to value at trial onset evident in our timecourse analysis, traditional whole-brain event-related GLM analyses confound online value responses with the offset of offline value responses. However, we report the traditional GLM analyses here for completeness. Group differences were the strongest in clock-aligned analyses of modulation by prior value, where SA displayed more negative value responses compared to SI in the left caudate nucleus and right putamen (ptfce $p_{FWE}$<0.05, whole brain, bi-sided; eTable 3), consistent with the timecourse findings that attempters had elevated striatal control network activity at low values. Additionally, at an exploratory threshold of p<0.005, uncorrected, we saw a wide-spread pattern of blunted value encoding in the attempters compared to comparison groups in other regions including the medial PFC, hippocampus, and bilateral striatum (eFigure 3A). Feedback-aligned analyses of updated value responses yielded similar but weaker results (eFigure 3B).

#### Region of interest (ROI) fMRI GLM coefficient results

We then conducted a more directed analysis of extracted $V_{max}$ fMRI coefficients for prior value (aligned to clock onset) and updated value (aligned to feedback) from anatomically defined regions of interest known to be implicated in value encoding (36–38) including the ventral prefrontal cortex (vPFC) and subcortical areas (putamen, caudate nucleus, nucleus accumbens, hippocampus, and thalamus). The regions were classified into networks (default mode, limbic, control, and hippocampus/amygdala) based on functional parcellations (29, 36, 39–41). We ran linear mixed effects models (lmer in R, version 4.1.0) for the vPFC and subcortical regions separately, with the extracted ROI values as the dependent variable and group, network, and group-by-network interaction as the fixed effects, with subject included as a random effect. Bonferroni FDR correction for multiple comparisons at a threshold of p<0.05 was used to determine significance, correcting for the four separate models that were tested.

Overall, we found that SA tended to have blunted $V_{max}$ response, with the strongest effects seen around the time of clock onset (prior value) and in the subcortical regions. Specifically, we found a significant group by network interaction for prior value in the subcortical areas (df = 9, eFigure 2C). Post-hoc pairwise comparisons (emmeans, p-values FDR-adjusted for six tests within each network) showed that SA had significantly lower value responses compared to D-NS in the default network (t=-2.92, p=0.022, df=457) and hippocampus/amygdala (t=-3.02, p=0.019, df=129). Additionally, SA had lower fMRI $V_{max}$ coefficients compared to SI in the control network (t=-3.38, p=0.004, df=457) and limbic network (t=-2.95, p=0.020, df=457). SA showed a trend towards lower fMRI $V_{max}$ coefficients compared to D-NS individuals in the control network (t=-2.014, p=0.089, df=457). Finally, there was also trend towards D-NS individuals having lower fMRI $V_{max}$ coefficients compared to C in the hippocampus/amygdala (t=-2.40, p=0.054, df=129). In the vPFC, the group by network interaction for prior value was not significant (eFigure 2A), but post-hoc pairwise comparisons showed that SA had lower fMRI $V_{max}$ coefficients compared to D-NS in the limbic network (t=-2.70, p=0.044, df=216), with a trend in the default network (t=-2.52, p=0.058, df=171). Additionally, SA showed a trend towards lower fMRI $V_{max}$ coefficients compared to SI in the default network (t=-2.36, p=0.058, df=171) and limbic network (t=-2.22, p=0.083, df=216). For updated value, there was a trend towards a group by network interaction in the vPFC (df=6, eFigure 2B), but none of the post-hoc pairwise comparisons were significant. Finally, the group by network interaction in the subcortical areas was not significant (eFigure 2D), but post-hoc pairwise comparisons revealed that SA had a trend towards lower fMRI $V_{max}$ coefficients in the control network compare to D-NS (t=-2.45, p=0.051, df=273) and SI (t=-2.40, p=0.051, df=273).

#### Timecourse fMRI analysis of reward and prediction error

In striatal subregions receiving projections from limbic and control cortical networks, high prior value predicted lower activity during the subsequent trial, consistent with reward prediction error dynamics (Figure 2A, 3A; see eFigure 4 for dynamics of BOLD response to reward and reward prediction error). In contrast, prior value responses persisted in the vPFC default network and the hippocampus. Altogether, as values of locations in the one-dimensional space (response times) were being learned, activity in the prefrontal default network best matched axiomatic value signals and striatal activity (except for subregions receiving default network projections) matched axiomatic reward prediction error signals. The prefrontal limbic network and amygdala displayed an intermediate pattern of activity, overall consistent with modulation by value signals, but without prior value persistence during the trial.
